# Compartments in medulloblastoma with extensive nodularity are connected through differentiation along the granular precursor lineage

**DOI:** 10.1101/2022.09.02.506321

**Authors:** David R. Ghasemi, Konstantin Okonechnikov, Anne Rademacher, Stephan Tirier, Kendra K. Maass, Hanna Schumacher, Julia Sundheimer, Britta Statz, Ahmet S. Rifaioglu, Katharina Bauer, Sabrina Schumacher, Michele Bortolomeazzi, Felice Giangaspero, Kati J. Ernst, Julio Saez-Rodriguez, David T. W. Jones, Daisuke Kawauchi, Jan-Philipp Mallm, Karsten Rippe, Andrey Korshunov, Stefan M. Pfister, Kristian W. Pajtler

## Abstract

Medulloblastoma with extensive nodularity (MBEN) are cerebellar tumors with two histologically distinct compartments and varying disease course. In some children MBEN progresses, while others show spontaneous differentiation into more benign tumors. However, the mechanisms that control the tug-of-war between proliferation and differentiation are not well understood. Here, we dissected this process with a multi-modal single cell transcriptome analysis. We found that the internodular MBEN compartment comprised proliferating early cerebellar granular neuronal precursors (CGNP)-like tumor cells as well as stromal, vascular, and immune cells. In contrast, the nodular compartment consisted of postmitotic, neuronally differentiated MBEN cells. Both compartments were connected through an intermediate cell stage of actively migrating CGNPs. Furthermore, astrocyte-like tumor cells were identified that had branched off the main CGNP developmental trajectory. Cells with an astroglial phenotype were found in close proximity to migrating, late CGNP-like and postmitotic neuronally differentiated cells. Our study reveals how the spatial tissue organization is linked to the developmental trajectory of proliferating tumor cells through a migrating precursor stage into differentiated tumor cells with a more benign phenotype. We anticipate that our framework for integrating single nucleus RNA-sequencing and spatial transcriptomics will help to uncover intercompartmental interactions also in other cancers with varying histology.

## Introduction

Medulloblastoma (MB) is the most common embryonal brain tumor of childhood and accounts for a significant proportion of both morbidity and mortality in this age group.^1, 2^ Traditionally, MB has been stratified into four histopathological subtypes based on histological appearance: Classic, Large Cell/Anaplastic, Desmoplastic/Nodular (DNMB) and MB with Extensive Nodularity (MBEN). ^3, 4^ Additionally, in the last decade, four molecular groups (WNT, SHH, Group 3, and Group 4) together with various subgroups have been defined and now generally replace the classic histopathological stratification in the 2021 WHO classification of central nervous system (CNS) tumors.^5–14^

MBEN is a unique histological type of MB that mainly arises in very young children and infants. These tumors exclusively fall into the molecular SHH-group and have been shown to frequently harbor mutations in SHH-pathway genes, for instance, *PTCH1* or *SUFU*.^4, 15^ Their distinctive, bicompartmental histological appearance is characteristic and differentiates them from other MBs: large nodular conglomerates of postmitotic, differentiated, neurocytic tumor cells are surrounded by internodular zones of highly proliferative, poorly differentiated cells.^15–17^ Furthermore, MBEN show a distinctive, grapevine-like neuroradiological appearance.^16–18^ MBEN are generally associated with good prognosis and are therefore considered low-risk in comparison to other MB variants. However, relapse and disease progression can occur and may require intensified therapy, resulting in potentially life-long, severe side effects in these very young children.^15, 19–22^ Interestingly, case studies reported that MBEN may mature and differentiate into benign ganglion cell tumors.^23, 24^ These findings have led to speculations whether underlying biological differentiation programs may exist in these tumors that could result in a loss of malignant potential.^16^ This hypothesis is underlined by the fact that MBEN are defined by the upregulation of cellular pathways that are important in cerebellar neuronal differentiation, for instance synaptic transmission, glutamatergic signaling and calcium homeostasis.^25^ In the past, it has been disputed whether DNMB and MBEN represent two biologically distinct MB variants. We previously showed that these histological variants can be reliably distinguished based on their transcriptomic profiles, despite harboring similar SHH-associated epigenetic signatures.^25^

Whereas clinical characteristics of MBEN are well established, the biology underlying its distinctive histological appearance and varying disease aggressiveness remains largely unknown. It is especially unclear if and how the two histological compartments of MBEN are interconnected. Several studies have postulated that the cells of origin of SHH-MB are cerebellar granule neuronal precursors (CGNP) in the external granular layer (EGL) of the developing cerebellum. The large differences in biological and clinical features within the molecular group of SHH-MB indicate that tumor formation may depend on distinctive spatial and temporal circumstances. ^26–30^ In this study, we used an integrated multi-modal approach that included three complementary spatial transcriptomics methods. We demonstrate that MBEN mimics the development of CGNPs into granule neurons and that the bi-compartmental histology of MBEN represents distinct differentiation cell states that are connected through a direct developmental trajectory. Furthermore, we identified a subset of tumor cells that cluster together with non-malignant astrocytes, show an astroglial phenotype, and branch off the main developmental trajectory in MBEN development. Overall, our findings indicate that MBEN could be understood as a disease of the developing cerebellum and serve as an explanation why these tumors are almost entirely restricted to infancy.

## Results

### Molecular and clinical characterization of the MBEN cohort

In order to study the genetic basis of MBEN in a comprehensive way, we applied a multimodal set of complementary methods, including single nucleus RNA-sequencing (snRNA-seq) and spatial transcriptomics, to a cohort of nine fresh frozen MBEN samples (**Fig. 1A, Suppl. Tbl. 1**). DNA methylation-based clustering of bulk methylomes with a reference cohort spanning all major molecular MB-groups confirmed that all cases belonged to the infant SHH-MB group (SHH-1: n = 7, SHH-2: n = 2) (**Fig. 1B**).^9^ DNA sequencing revealed genetic alterations of the SHH-pathway in six tumors (*PTCH1*: 1/9, *SMO:* 2/9, *SUFU:* 3/9)(**Fig. 1C**). As reported previously for MBENs, copy number variations (CNVs) were only detected infrequently (**Suppl. Fig. 1**). MBEN-histology was confirmed by central pathological review in all cases (A.K.) (**Fig. 1D**). All tumors were located in the cerebellum (**Fig. 1E**). At the time of diagnosis, disease stage was M0 in 7/9 cases, with two patients showing metastatic disease (M2: 1/9, M3: 1/9) (**Suppl. Fig. 2A**). The median age of children included in this cohort was 2 ± 0.70 years and 7/9 patients were male (**Suppl. Fig. 2B, C**). In accordance with earlier studies, the clinical outcome in the presented cohort were overall favorable. However, a total of three out of nine children experienced relapses of their disease (**Suppl. Fig. 2D, E**).

**Fig. 1.**
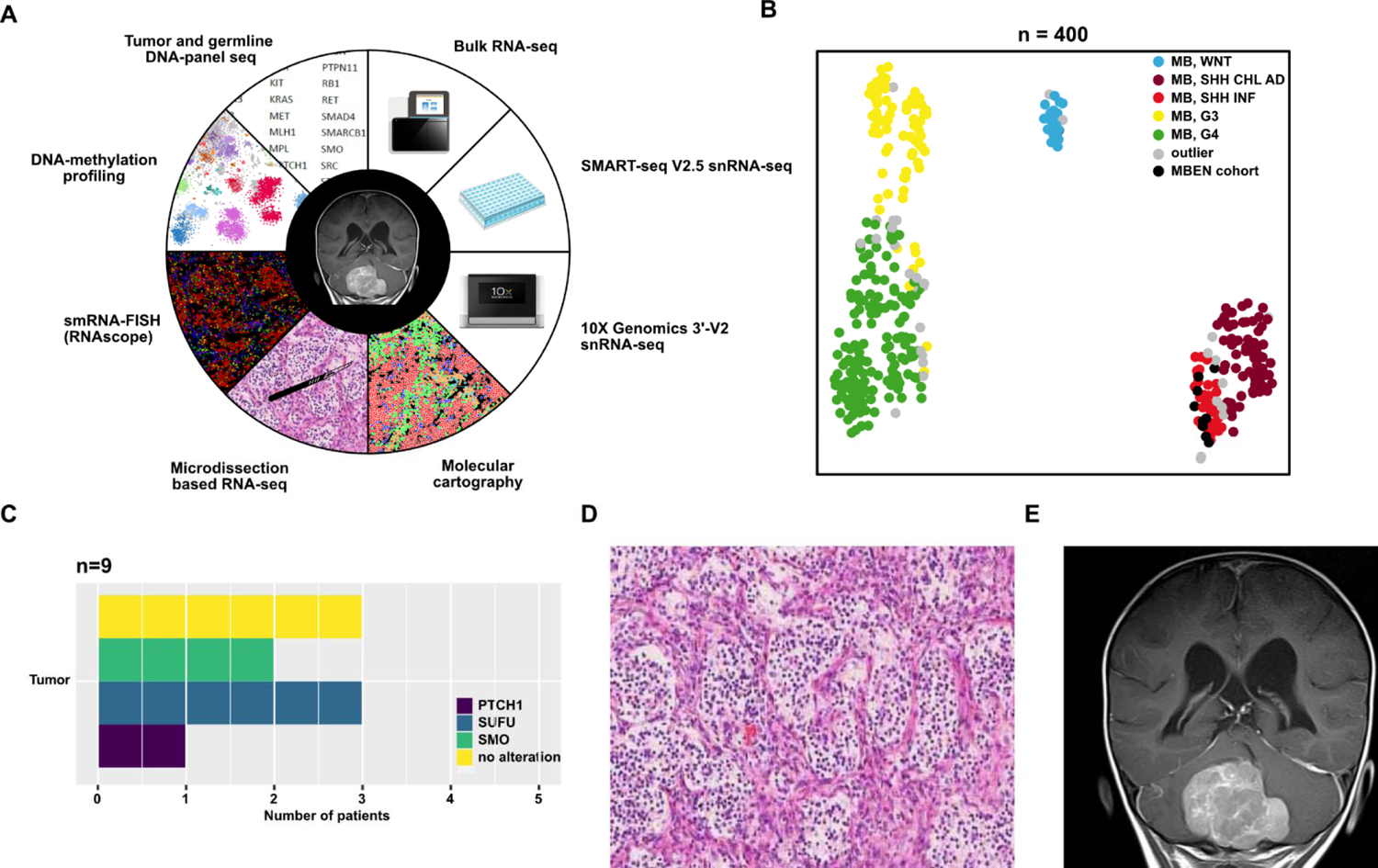
Method spectrum and molecular features of the MBEN patient cohort. **A** Visual summary of all methods that were applied to investigate the MBEN cohort. **B** t-SNE depicting a clustering of the MBEN cohort (n = 9) with a reference cohort of 391 molecularly characterized MBs. All nine samples clustered with the SHH-subtype. **C** Bar chart summarizing the alterations in the SHH-pathway in both blood and tumor that were detected using DNA panel seq. **D** Representative H.E. staining showing the two characteristic histological components of MBEN. **E** Representative coronal MRI image in an MBEN patient showing a large cerebellar mass with grapevine-like appearance.

### A subset of tumor cells is transcriptionally similar to non-malignant astrocytes

To investigate intra- and intertumoral transcriptional heterogeneity within MBEN, we isolated nuclei from fresh frozen material and subsequently applied two complementary methods of snRNA-seq (see methods), namely the 10X V2 3’- (n = 9) and the SMARTseq V2.5-protocols (n = 6). After initial quality control, 28132 and 1526 nuclei were used for downstream analysis, respectively. In order to ensure that snRNA-seq datasets faithfully represented the respective tumor, a fingerprint analysis using bulk RNA-sequencing profiles was conducted (**Suppl. Fig. 3A**). Furthermore, comparative clustering revealed concordant results between both methods, confirming that both techniques covered the same cell types (**Suppl. Fig. 3B**). Subsequently, we integrated both datasets to arrive at an integrated snRNAseq dataset resulting in 29658 cells for further in-depth analysis (**Fig. 2A, Suppl. Fig. 3C, D**).^31^ Non-malignant cells were identified based on established marker genes or recently published single cell atlases and comparison to a reference dataset of the fetal cerebellum, which we recently published.^32–34^ This approach resulted in the identification of astrocytes, oligodendrocytes/oligodendral precursors, microglia/ endothelial and fibroblast/ perivascular cells (**Fig. 2B, Suppl. Fig. 3E, F**). Apart from microglia, no significant immunological infiltration, for instance by T- or NK-cells, was observed, confirming earlier studies that described MB as an immunologically cold tumor.^35^ Surprisingly, analyzing genome-wide copy number variations (CNV) per individual cluster using inferCNV revealed a CNV-signature in the astrocytic cluster (**Fig. 2C**).^36^ In order to identify which cells in this cluster were malignant and non-malignant, respectively, we analyzed both snRNA-seq datasets separately and without correcting for patient-related batch effects. We hypothesized that malignant cells would cluster according to patient, while non-malignant cells would congregate in one cluster, as observed in other single cell studies (**Suppl. Fig. 3E, F**).^37^ Using this approach, we could indeed separate malignant from non-malignant cells within the astrocytic cluster, which partly clustered in mixed, non-malignant, and partly in malignant, patient-specific clusters (**Fig. 2D – G**). Whereas the number of tumor astrocytic cells varied by sample, they were identified in every case (**Suppl. Fig. 3G**). Based on these findings, we hypothesized that the subpopulation of MBEN cells with astrocytic features represent tumor astrocytes, which is supported by experimental studies that indicate that murine MB cells can transdifferentiate into tumor-derived astrocytes.^38–40^

**Fig. 2.**
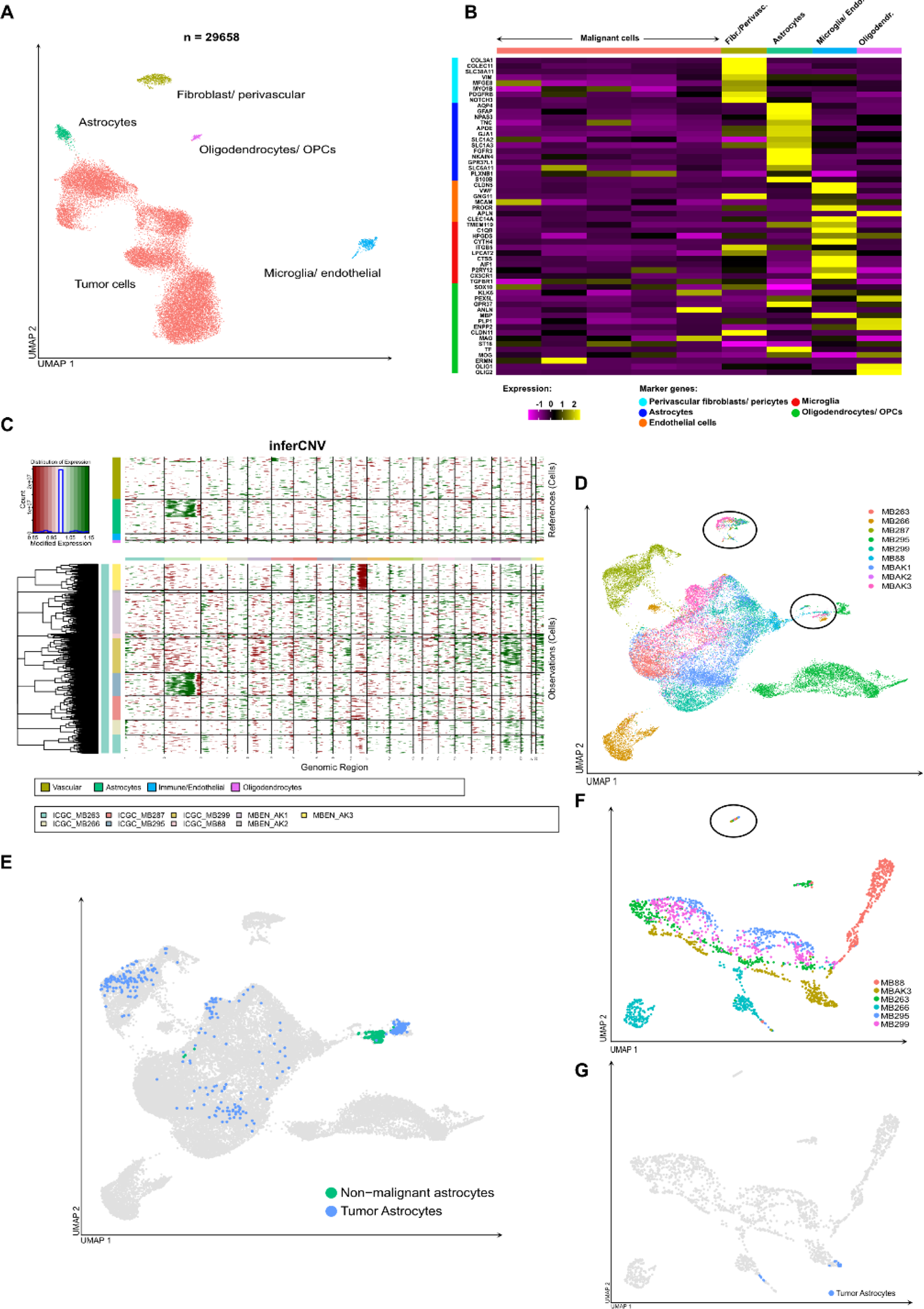
SnRNA-seq reveals a subset of MBEN-cells with similarity to astrocytes. **A** UMAP-projection of the integrated dataset (10X snRNA-seq + SMARTseq V2.5 snRNA-seq) shows intratumoral heterogeneity in MBEN. **B** Heatmap depicting marker genes for indicated cell types of all nine clusters. The first five clusters represent malignant MBEN cells. Clusters 6 – 9 are identified as perivascular fibroblasts/ pericytes, astrocytes, oligodendrocytes/ oligodendroglial precursors and a mixture of microglia and endothelial cells. **E** InferCNV-analysis using non-malignant cells as a control confirms malignant origin for the vast majority of cells as well as a CNV signature of malignant cells in the astrocyte cluster. **D** UMAP projection of the 10X snRNA-seq dataset which was not corrected for patient-associated batch effects. Whereas the majority of cells are clustering according to patient, significant mixing of cells occurs in two clusters (encircled), representing non-malignant cells. **E** UMAP projection as in Fig. 2D (10X snRNA-seq dataset) in which non-malignant and tumor astrocytes are highlighted. Whereas a fraction of the cells from the astrocyte cluster (compare Fig. 2A) falls into one of the two mixed, non-malignant clusters, the other part clusters with malignant MBEN cells. **F** UMAP projection of the SMARTseq V2.5 snRNA-seq. dataset which was not corrected for patient batch effect. Whereas the majority of cells cluster according to patient, significant mixing of cells occurs in one cluster (encircled), representing non-malignant cells. **G** UMAP projection (SMARTseq V2.5 dataset) in which tumor astrocytes are highlighted. No cells from of the astrocyte cluster (compare Fig. 2A) falls into the mixed, non-malignant cluster.

Taken together, we found that besides malignant cell subpopulations MBEN contains various non-malignant cells with both glial and mesenchymal transcriptional signatures. Interestingly, snRNA-seq using two complementary methods revealed a distinct subpopulation of tumor cells that correspond to tumor astrocytes.

### Cell stages in MBEN recapitulate cerebellar granular development

In order to study MBEN cells in greater detail, non-malignant cell populations, including non-malignant astrocytes, were excluded from further tumor cell analysis. Re-analysis restricted to malignant cells revealed five distinct clusters and one additional one, which was entirely driven by upregulation of heat shock- and ribosomal-associated genes, indicating an artificial effect most likely induced by cellular stress associated with sampling (**Fig. 3A**). Only one cluster was actively proliferating as indicated by S- and G2M-signatures (**Fig. 3B**). The different cell populations showed distinct expression patterns matching those of individual CGNP developmental stages. Two clusters, of which one was proliferating, showed upregulation of marker genes of early CGNPs in the EGL (e.g., *BARHLH1, ZIC1*, *ZIC3, PTPRK*) and SHH-pathway members (*PTCH1, SMO, HHIP, GLI1, GLI2)*. Additionally, the proliferating cluster expressed genes that are implicated in epigenetic regulation and chromatin remodeling (e.g., *EZH2, SUZ12, CTCF, YY1, RAD21*)(**Fig. 3C, Suppl. Fig. 4, Suppl. Tbl. 2**).^41–45^ These genes have been shown to be involved both in the orchestration of cerebellar development and the oncogenesis of numerous cancer types.^45, 46^ Clusters three and four were characterized by the expression of markers of intermediately differentiated, migrating (*GRIN2B, CNTN2, ASTN1, SEMA6A*) and postmitotic, differentiated CGNPs (*GABRA1, GABRA6, GRIN2C*), respectively (**Fig. 3C, Suppl. Fig. 4**).^41, 47–49^ A fifth cluster was formed by tumor astrocytes. Notably, we observed that markers associated with astrocyte-immune cell interactions in murine models (e.g. *IGF1, IL4*) did not show any specificity in human tumors, being spread across all cell types in MBEN samples.^38^ In addition, tumor astrocytes showed upregulation of stromal and of early cell stage markers (*LAMA2*, *SOX2*, *SOX9*), but no significant proliferative activity. Pseudotemporal ordering revealed a continuous lineage starting from the proliferating, early CGNP-like cell states, spanning the intermediate ones and finally congregating in the postmitotic, neuronally differentiated cells (**Fig. 3D**). This lineage resembled physiologic differentiation of CGNPs into normal granular neurons during cerebellar development. Interestingly, tumor astrocytes branched off early in the trajectory and prior to the appearance of markers of cell migration (**Fig. 3D**). These results corroborated an earlier study indicating that tumor astrocytes are not tumor stem cells, which was further supported by the fact that the respective cluster did not show significant proliferative activity.^38^ Correlating our snRNA-seq data with transcriptomic profiles of the aforementioned study, MBEN-derived tumor astrocytes showed clear similarity to both normal and tumor astrocytes from the respective SHH-MB mouse model (**Fig. 3E**). In order to further validate our findings, we mapped our MBEN dataset onto a comprehensive snRNA-seq atlas that we recently published, spanning the whole embryonal and fetal development of the cerebellum^34^. This analysis confirmed results of the pseudotime trajectory analysis. Whereas cells from the beginning of the MBEN trajectory resembled transcriptional signatures of early CGNPs, the intermediate MBEN cluster showed the highest concordance with intermediate developmental stages of CGNP differentiation (**Fig. 3F**). Lastly, the neuronal-like cluster at the end of the trajectory clearly mapped onto differentiated granular neurons and tumor astrocytes showed high similarity with astrocytes from the developing cerebellum.

**Fig. 3.**
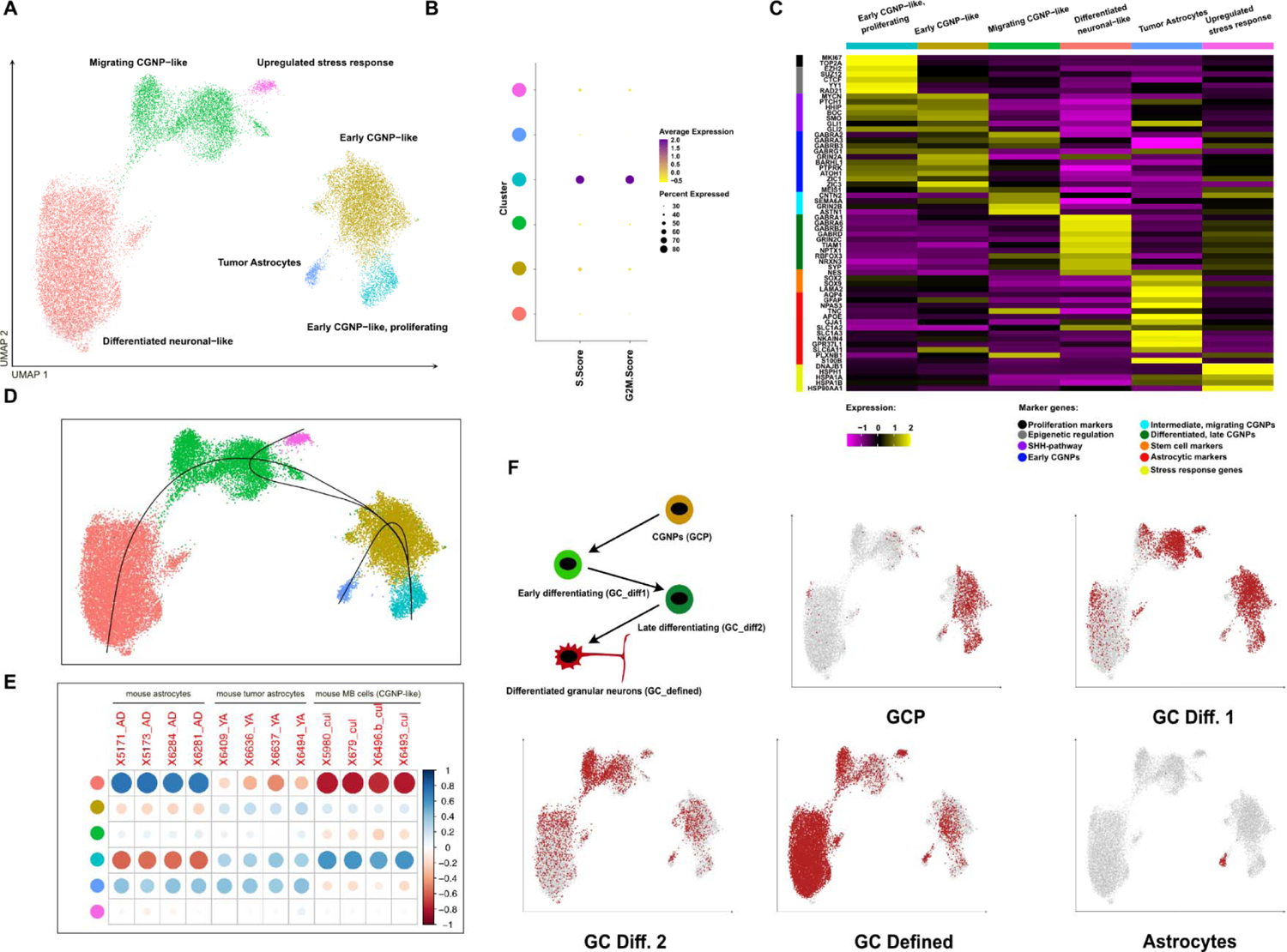
MBEN differentiates along a cerebellar developmental trajectory. **A** UMAP-plot of the merged snRNA-seq-dataset (10X snRNA-seq + SMARTseq V2.5 snRNA-seq) restricted to malignant cells only. Clusters are named based on their similarity to major stages of physiological cerebellar granular neuronal precursors (CGNP) development. **B** Dotplot illustrating proliferation activity based on gene signatures of S- and G2M-cell cycle activity, which is restricted to one cluster only. Color code of the y-axis as in A. **C** Heatmap showing the expression of important cerebellar developmental genes in the dataset per indicated cluster. **D** UMAP-projection with overlayed pseudotime trajectories shows two major lineages, with the majority of cells following a granular cerebellar trajectory, and a small subset differentiating into tumor astrocytes. **E** Tumor astrocytes show transcriptional similarity to murine astrocytes and murine tumor astrocytes, but not to MB cells with transcriptional similarity to CGNPs (Yao *et al.*, Cell, 2020). Color code of the y-axis as in A. **F** Feature plots of the dataset highlighting similarity of MBEN associated tumor cells to non-malignant cells of the developing cerebellum. GCP = granular cerebellar precursors (Okonechnikov *et al.*, bioRxiv, 2021).

In summary, different MBEN cell populations mimicked distinct cell stages of the normal CGNP lineage, spanning the whole developmental trajectory. The cell stages of MBEN development differed markedly in their major expression programs, and proliferative activity was lost after the early CGNP-like MBEN cells started to differentiate. Interestingly, tumor astrocytes branched off early from the main developmental trajectory within MBENs.

### Functional *in silico* analyses reveal biological mechanisms underlying MBEN differentiation

A number of core biological mechanisms and cellular functions play a pivotal role in CGNP development. These include, amongst others, the SHH pathway, NMDA/glutamate signaling, the expression of bone morphogenic proteins (BMPs) as well as Ca^2+^-signaling.^41, 42, 47, 48, 50–55^ We sought to investigate whether similar processes might influence MBEN development. First, we examined intra- and intercluster communication by analyzing ligand-receptor pairs based on snRNA-seq expression data (**Fig. 4A, B**).^56^ Tumor astrocytes were the only cells that showed high-confidence interactions with all other clusters via *APOE* signaling, which is in line with previous findings that astrocytic cells act as the main supplier and redistributor of cholesterol.^57^ Intra-cluster communication within the differentiated neuronal-like cluster was dominated by genes which are involved in voltage-dependent signaling, especially with regard to Ca^2+^-signaling, such as *RIMS1, CACNA1C* and *CALM1*.^58^ These findings were corroborated by analyzing the expression of a transcriptomic Ca^2+^-signaling signature including 1805 genes.^59^ Indeed, Ca^2+^-signaling was strongly connected to later cell stages within MBEN (**Fig. 4C**). Another group of genes that is involved in CGNP development relates to BMPs that may also suppress MB proliferation *in vitro*.^41, 60^ Interestingly, a BMP-related signature showed strong expression in tumor astrocytes, suggesting that these cells might be involved in triggering the differentiation process by suppressing proliferative activity in MBEN (**Fig. 4D**). The hypothesis that similar processes as in physiological CGNP development are active in MBEN was further confirmed by cluster-specific gene ontology (GO) analysis (**Suppl. Fig. 5, Suppl. Tbl. 3**).^61^ All clusters showed significant fold enrichments for GO terms directly connected to synaptic organization and activation, glutamate signaling, and Ca^2+^-homeostasis (**Suppl. Fig. 5, Suppl. Tbl. 3**). Tumor astrocytes showed high expression of genes connected to cAMP-metabolism, possibly indicating that malignant astrocytes may be involved in energy supply for surrounding cells, thus resembling astrocytic functions in normal CNS.^62^

**Fig. 4.**
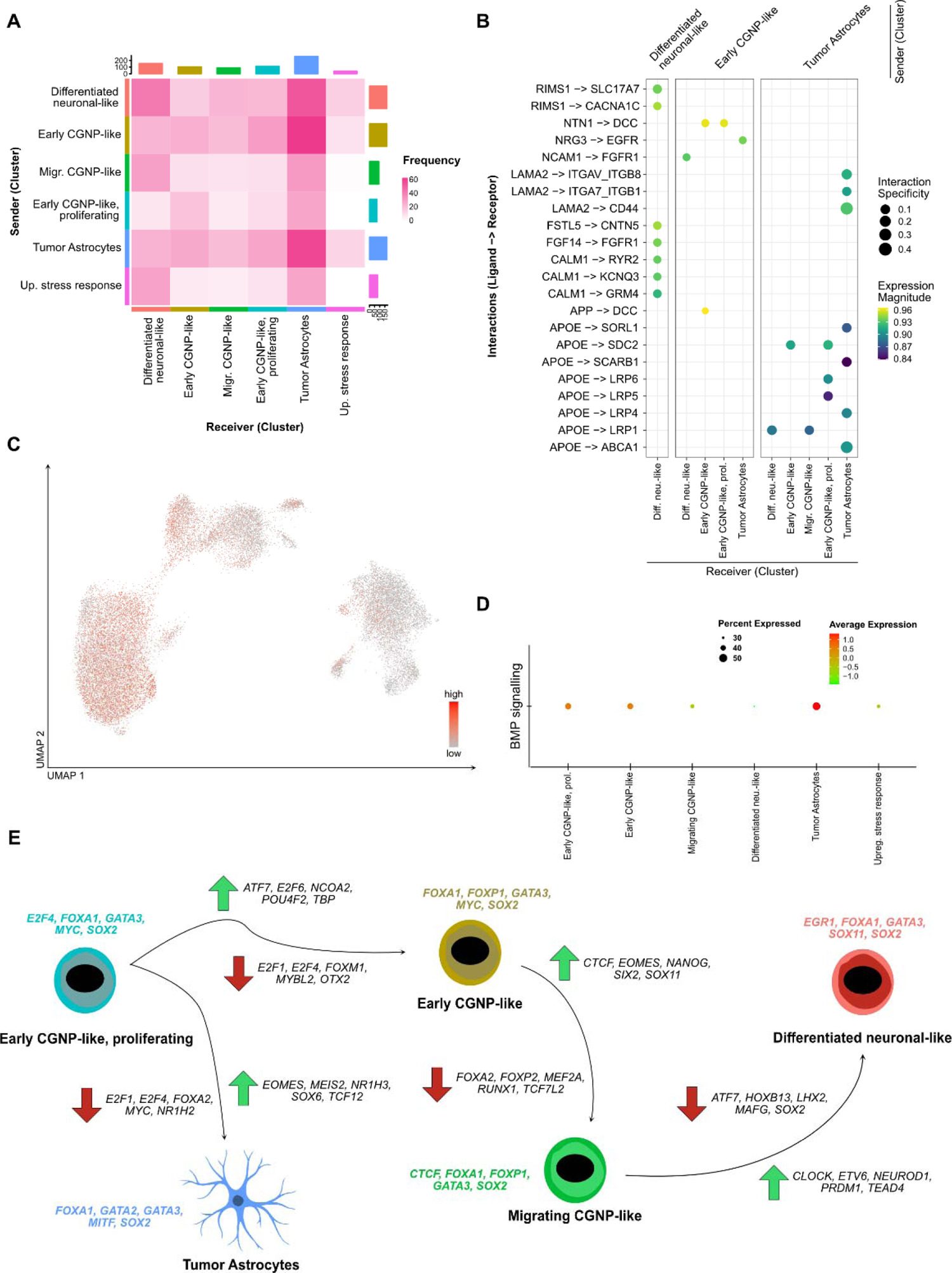
Biological processes that regulate MBEN differentiation mimic CGNP development. **A** Heatmap depicting the frequencies of predicted interactions for each pair of potentially communicating cell stages amongst MBEN cells. **B** Dotplots depicting the most confident receptor-ligand-interactions within and between different MBEN cell stages. The interaction specificity weights indicate how specific a given interaction is to the sender and receiver cell stages. **C** UMAP-plot showing the expression of a Ca^2+^ signaling signature along MBEN differentiation. To improve visibility, only cells with expression levels above the 25^th^ quantile are highlighted. **D** Dotplot showing the expression of a BMP protein signature along MBEN differentiation. **E** Visualization of transcription factor (TF) activities within and between MBEN cell stages. Above each cell stages, the top five expressed TFs are indicated. Furthermore, the five most up- and downregulated TFs are given for each developmental step of the MBEN trajectory (see also Fig. 3D).

In order to complement these findings, we performed cluster specific transcription factor (TF) activity analysis using the DoRothEA tool.^63, 64^ To this end, overall TF activity per cluster and changes in TF activity between MBEN stages that followed each other were calculated (**Fig. 4E, Suppl. Fig. 6, Suppl. Tbl 4, 5**). *SOX2* and *FOXA1*, two pioneer TFs which are involved in maintaining chromatin accessibility throughout early development^65^, were active along the entire differentiation process. In contrast, *E2F1* and *E2F4*, which are particularly involved in cell cycle regulation^66^, were downregulated shortly after the start of the trajectory. In addition, *MYC*, which was among the most active TFs in (proliferating) early-like CGNPs, was downregulated in later MBEN cell stages. *EOMES*, a marker of unipolar brush cells and intermediate cortical neurons^34, 67^, increased in activity once (proliferating) early CGNP-like cells differentiated into migrating CGNP-like cells and tumor astrocytes. Similarly, *NEUROD1*, a TF that has been described as an inducer of differentiation and thus tumor suppressor gene in MB, increased activity throughout the MBEN differentiation process (**Fig. 4E, Suppl. Tbl. 4, 5**). Surprisingly, despite clear differences in expression profiles, tumor astrocytes showed overall similar TF activities to other clusters, possibly due to their shared developmental roots. Notably, *NR1H3*, a TF that is strongly involved in cholesterol homeostasis^68^, showed markedly increased activity in tumor astrocytes.

Taken together, complimentary functional *in silico* analyses confirmed that biological mechanisms of physiological CGNP development are mimicked in MBEN differentiation. Furthermore, tumor astrocytes were connected to energy and cholesterol homeostasis.

### Transcriptomic analysis of MBEN based on microdissected tissue shows differences between the internodular- and nodular compartments

In order to correlate our findings with the bicompartmental histology of MBEN, we applied targeted microdissection to FFPE samples overlapping with our study cohort, allowing to obtain bulk RNA-seq datasets from nodular and internodular tumor areas separately (n=26). After expression variance inspection and fingerprint verification of corresponding samples (**Suppl. Fig. 7A,B**) we corrected for patient-related batch effects of the original tissue (details in methods) and detected genes differentially expressed between nodular and internodular compartments (**Fig. 5A, Suppl. Tbl. 6**). When we investigated the expression of these genes in our snRNA-seq datasets, we observed clear patterns. For example, *TRIM9*, which was nodular-specific in the microdissection data, was strongly expressed in intermediate CGNP-like and neuronal-like snRNA-seq clusters (**Fig. 5B**). Similarly, *TMEM108* from internodular derived microdissections demonstrated a strong fit to clusters of early CGNP-like MBEN cells and tumor astrocytes (**Fig. 5C**). To further confirm these findings, we performed gene set variation analysis^69^ that demonstrated differentiated MBEN clusters being strongly enriched with nodular, whereas the early CGNP-like populations were in close match to internodular transcriptome profiles (**Fig. 5D**). Interestingly, tumor astrocytes were more specific to the internodular compartment.

**Fig. 5.**
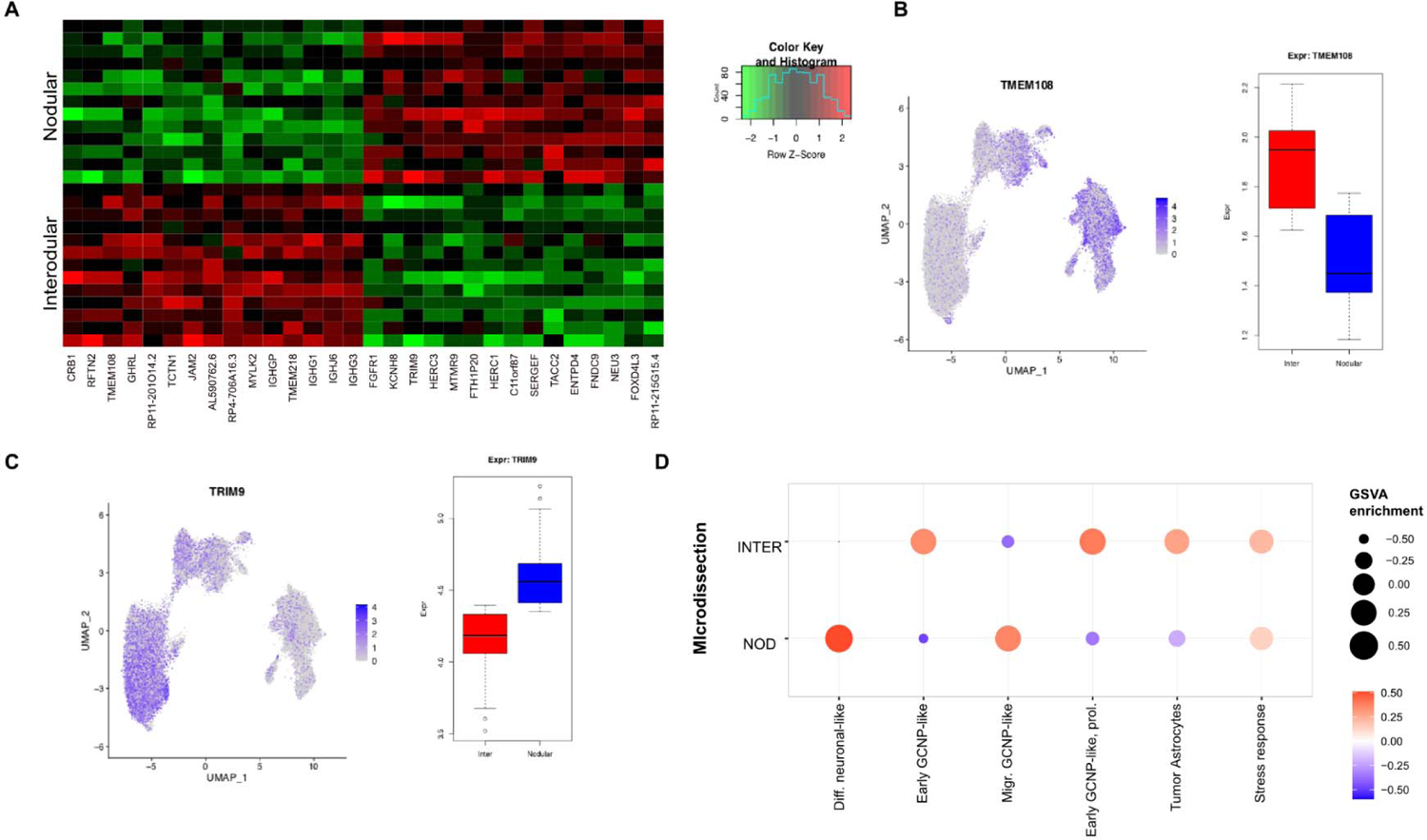
Microdissection reveals transcriptional differences between internodular and nodular tumor areas that correspond to distinct MBEN cell stages. **A** Heatmap showing differential gene expression between internodular and nodular MBEN compartments. **B** *TMEM108*, a marker of the internodular compartment, is mainly expressed in early CGNP-like cells. Feature plot on the left depicting gene expression in the snRNA-seq dataset. Bar chart on the right showing expression (measured in batch-effect adjusted RPKM) in microdissected MBEN tissue. **C** *TRIM9*, a marker of the nodular compartment, is mainly expressed in later stages of MBEN development. Feature plot on the left depicting gene expression in the snRNA-seq dataset. Bar chart on the right showing expression in microdissected MBEN tissue. **D** Gene set variance analysis (GSVA) enrichment of the expression signatures of microdissected internodular and nodular histological compartments to the different snRNA-seq clusters representing cell stages of the MBEN trajectory shows distinct transcriptional similarities between differentiated cells and the nodular compartment, and early-CGNP like cells with the internodular areas.

### Spatial transcriptomics correlates cell stages in MBEN to its histologic compartments

Our microdissection-based expression analysis revealed clear differences between the two MBEN compartments, however, it did not allow to investigate the detailed spatial distribution of the MBEN cell stages that were identified using snRNA-seq. Thus, we performed spatial transcriptomic analysis via single molecule RNA-*in situ* hybridization on three representative samples (smRNA-FISH/RNAscope). For RNAscope, twelve genes were chosen as markers for the different snRNA-seq clusters, and also considering biological significance (e. g. *GLI1* and *PTCH1* as SHH-pathway members) (**Fig. 6A, B, Suppl. Tbl. 7**). Two genes were excluded due to patient-specific expression or lack of discriminatory power in the spatial dataset, with 10/12 genes selected for downstream analysis (**Suppl. Fig. 8, 9**). After spot detection and nuclei segmentation, transcripts per nucleus were quantified and used to construct spatially derived single cells, which were then mapped back to the scanned image (**Suppl. Fig. 8, 9**). Based on marker gene expression alone, the bicompartmental histology of MBEN could be readily reconstructed. Using *RBFOX3*, a gene that encodes for the neuronal marker protein NeuN being specific for differentiated neuronal cells, and the embryonal gene *LAMA2,* found to be differentially expressed in non-malignant stromal cells and (tumor) astrocytes, we could distinguish between the nodular (*RBFOX3* high, *LAMA2* low), and internodular (*RBFOX3* low, *LAMA2* high) compartment (**Suppl. Fig. 10A - F**). These findings were further confirmed when we used the full set of all ten genes. Clusters of the RNAscope-derived single cells fully reconstructed both the snRNA-seq and microdissected tissue-derived findings, i.e., premigratory cycling and non-cycling early CGNP-like clusters, *LAMA2-*positive cells that included both stromal and astrocytic cells and migrating as well as postmigratory neuronally differentiated cells (**Fig. 6C**). Whereas the first three cell types were restricted to the internodular tumor compartment, postmigratory, differentiated cells were only found in the nodular areas (**Fig. 6D – G**). Intermediate, migrating cells were observed in both compartments. These visual observations were supported by quantifying the probability of each of the five cell types to be located next to each other, respectively, which confirmed that non-proliferating and proliferating early CGNP-like and *LAMA2*-positive cells (including astrocytic, vascular, and stromal cells) were co-localizing. In contrast, early CGNP-like and late stage neuronally differentiated MBEN cells were clearly less likely to be located next to each other (**Fig. 6H – J**).

**Fig. 6.**
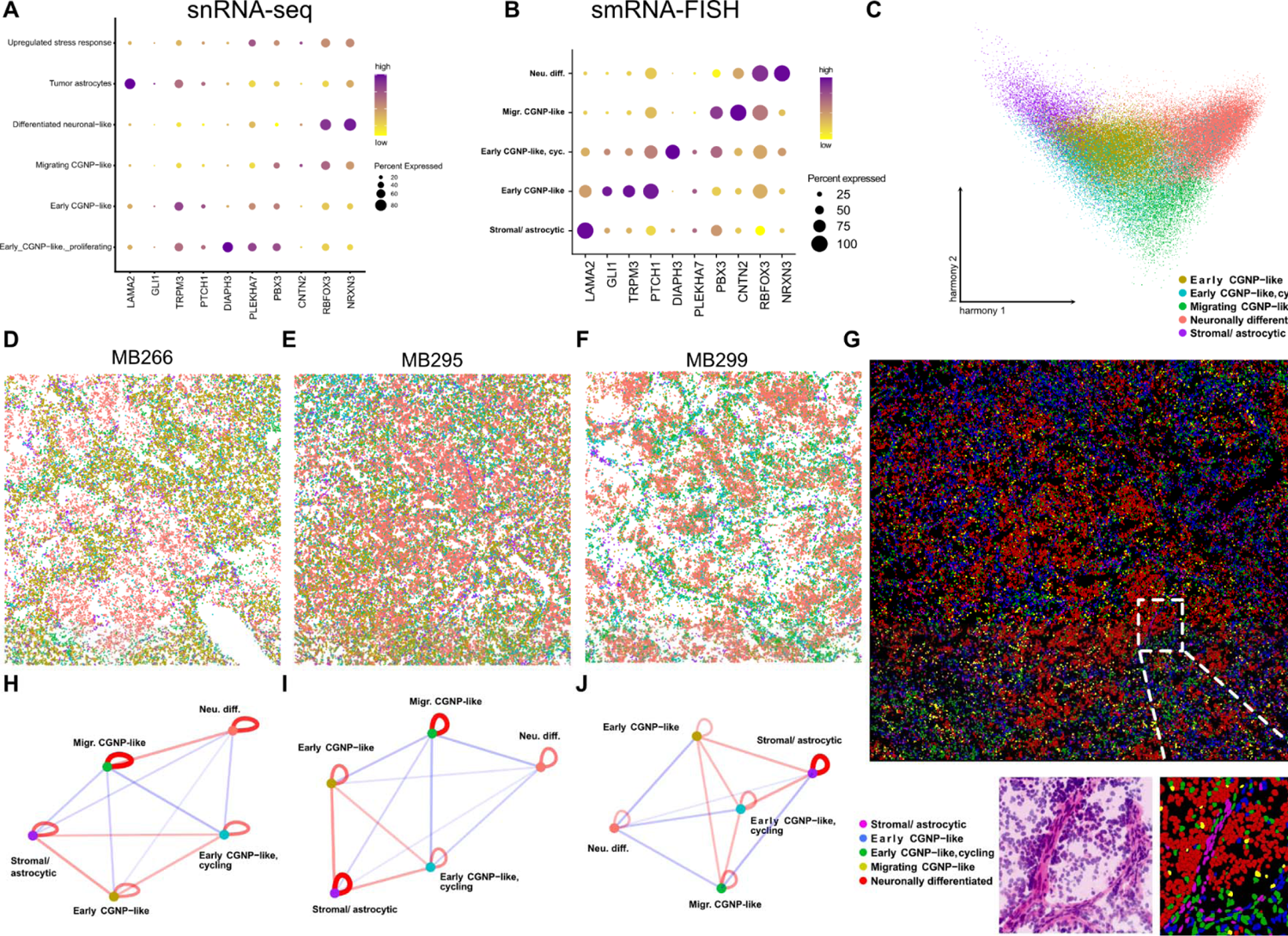
Spatial transcriptomics map developmentally distinct cell stages to spatially distinct tumor compartments. **A, B** Dotplots showing the expression of the ten genes chosen for spatial transcriptomics in the snRNAseq-**(A)** and the smRNA-FISH-dataset **(B)** for each cluster, respectively. **C** Harmony corrected UMAP-clustering of smRNA-FISH-derived single cells shows distinct clusters **D – F** Mapping single cells back to the original scans reveals spatially distinct localization of cells from each cluster with regard to the bicompartmental structure of MBEN. Color code corresponds to C. **G** Representative section of MB295 shows clear correlation of the smRNA-FISH-derived clusters to spatially distinct regions of MBEN. Lower panel left: Zoom-in with pseudo HE staining. Lower panel right: Corresponding mapping of single cells illustrates the spatial architecture of the transition zone between the two histological compartments of MBEN, which is mainly formed by early CGNP-like, migrating and stromal/astrocytic cells. **H – I** Network visualizations quantifying the probability of co-localization of cells from each cluster. Red lines and blue lines indicate high and low probability for co-localization, respectively. (Proliferating) early CGNP-like and stromal/astrocytic cells are co-localizing, whereas early and late stages of the MBEN trajectory are spatially separated. Color code corresponds to C.

Taken together, a targeted spatial transcriptomics approach confirmed our observations based on snRNA-seq and revealed that different normal CGNP-related cell stages of the MBEN developmental lineage could be mapped to the internodular and nodular compartments, respectively.

### The tumor microenvironment in MBEN differs between internodular and nodular compartments

The restricted number of marker genes using smRNA-FISH did not allow to differentiate between astrocytic and stromal cells. We therefore extended the spatial analysis by using the Molecular Cartography platform from Resolve Biosciences on four representative samples. In line with our prior findings, the expression patterns of single genes, such as *LAMA2* and *NRXN3*, were already sufficient to reconstruct the bicompartmental MBEN histology on the transcriptomic level (**Fig. 7A, Suppl. Fig. 11A - D**). Notably, the gene *TMEM108* (identified as differentially expressed in our microdissection-based gene enrichment analysis) could be confirmed as a marker for the internodular MBEN compartment (**Suppl. Fig. 11B**). In total, 92666 DAPI-segmented cells were subjected to downstream analysis. We were able to recover all major MBEN cell stages, namely proliferating and non-proliferating early CGNP-like, migrating CGNP-like and neuronally differentiated tumor cells (**Fig. 7B, C, Suppl. Tbl. 8**). Furthermore, one small cluster was dominated by proliferative activity only, whereas another cluster expressed genes of CGNPs in later developmental stages and was thus termed “late CGNP-like”. These two clusters, which were not distinguished as clearly in our snRNA-seq data, most likely represented subsets of proliferating, early CGNP-like and late stage neuronally differentiated MBEN cells. In addition, the designed extended gene panel allowed for identification of non-malignant cell types, such as stromal, vascular/endothelial, immune cells and (tumor) astrocytes (**Fig. 7B, C, D**). The immune cell cluster mainly consisted of a mix of microglia and bone marrow-derived macrophages (**Fig. 7C**). Interestingly, oligodendrocytes seemed to be generally underrepresented in the MBEN tumor microenvironment. Non-malignant astrocytes could not be distinguished from tumor astrocytes due to missing CNV information. However, we observed that a fraction of tumor cells was transcriptionally similar to MBEN tumor cells (**Figure 7D**). Notably, astrocyte marker genes, such as *NPAS3*, *CD44* and *LAMA2*, were also expressed in a fraction of early CGNP-like cells, which were identified as potential precursors of tumor astrocytes in our snRNA-seq trajectory analysis (**Fig. 3E, 7B,E, Suppl. Fig. 11A,C,D**).

**Fig. 7.**
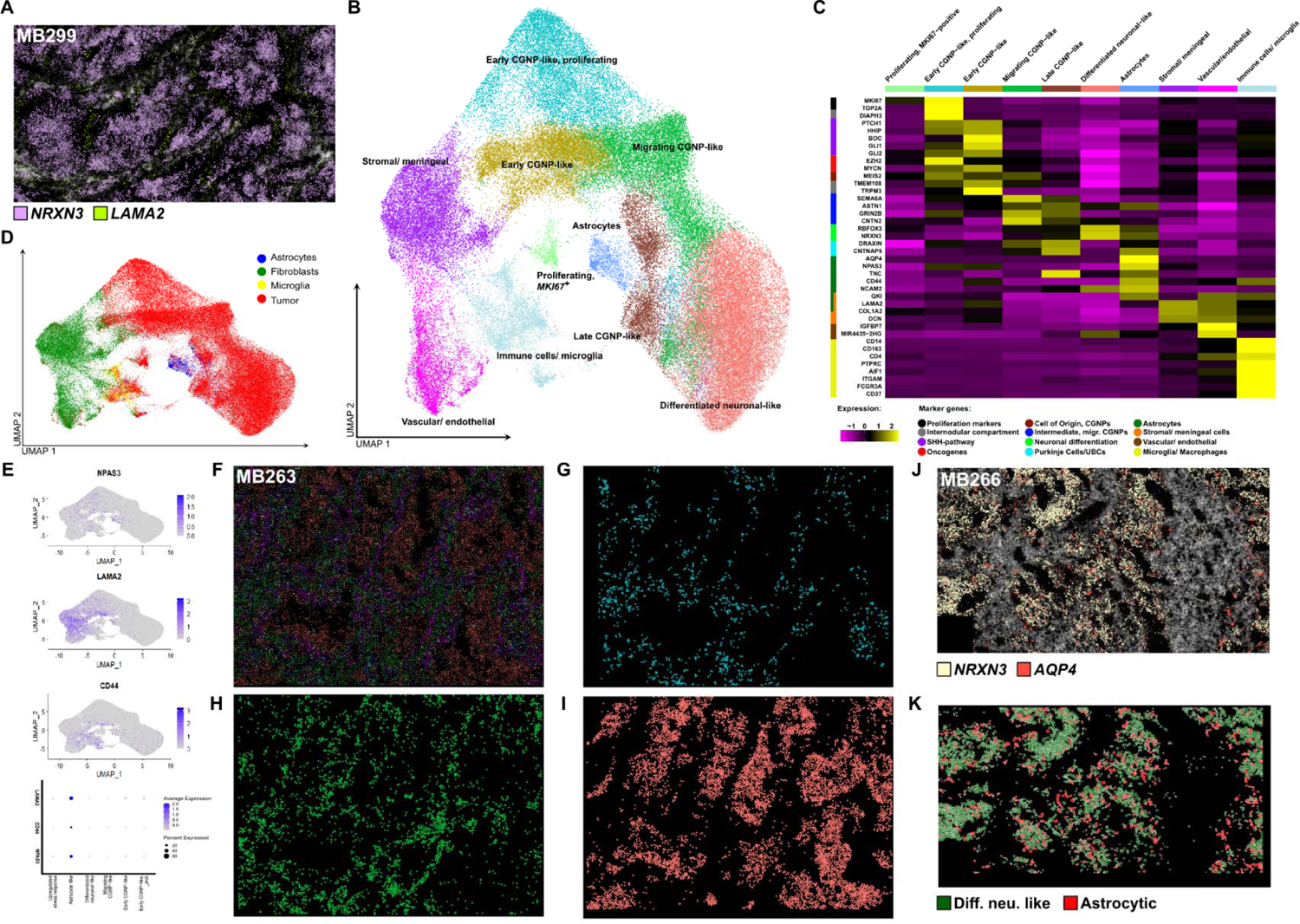
The distribution of non-CGNP-like cells in MBEN differs between internodular and nodular tumor compartments. **A** Representative scan of MB299 shows gene expression of the neuronal differentiation marker *NRXN3* (purple) and *LAMA2*, a marker gene that is strongly expressed in stromal cells and (tumor) astrocytes (green). The spatial expression of these two genes is sufficient to reconstruct the bicompartmental structure of MBEN. **B** UMAP projection showing the clustering of single cells derived from molecular cartography, which reconstructs all stages of MBEN development that were identified with snRNA-seq alongside none-malignant cell types. **C** Heatmap depicting marker genes of the different clusters from C. **D** Single cells were mapped onto the full MBEN snRNA-seq dataset, with clear evidence that non-malignant cell types could be assigned to astrocytes, fibroblasts, and microglia, respectively. **E** UMAP-plots showing the expression of the astrocytic marker genes *NPAS3, CD44* and *LAMA2*, which are expressed in astrocytic and a fraction of early CGNP-like cells. Bar chart shows the respective expression in MBEN clusters from snRNA-seq data. **F** Representative scan of MB266 in which single cells were projected back into the spatial space. By integrating all cell types derived from single cell construction, the bicompartmental structure of MBEN is reconstructed. The color code is equivalent to B. **G - I** Mapping of **(G)** proliferating, early CGNP-like, **(H)** intermediate, migrating, and **(I)** neuronally differentiated MBEN-cells. Whereas early CGNP-like cells are restricted to the internodular compartment, neuronally differentiated cells form the tumor nodules. Migrating MBEN-cells are spread throughout both compartments. **J** Expression patterns of *NRXN3* and *AQP4* (astrocyte markers) in a representative scan of MB266 **K** Astrocytic cells (red) are located at the transition border of the nodular compartment, which is formed by differentiated, neuronal-like cells (green).

Next, we sought to investigate the spatial distribution of each cell type based on the annotation described above. Similar to the results that we observed using smRNA-FISH, the internodular compartment was formed by (proliferating) early GCNP-like cells, whereas the nodular compartment consisted of neuronally differentiated tumor cells. Intermediate cell stages that expressed markers of migrating CGNPs were found in both histological areas (**Fig. 7F – I**). We then used cell proximity network analysis to investigate which cell types co-localize in MBEN (**Suppl. Fig. 11E - H**). Strikingly, there were marked differences between astrocytes/tumor astrocytes and other non-malignant cells in terms of spatial distribution (**Suppl. Fig. 11E – H, Suppl. Fig. 12A - T**). Stromal cells, vascular cells, and immune cells, which clustered separately from tumor cells in the UMAP-projection, were strongly enriched in the internodular compartment and co-localized with early CGNP-like, proliferating MBEN cells. In contrast, cells with an astroglial phenotype were found in close proximity to migrating, late CGNP-like and postmitotic neuronally differentiated cells. However, the majority of astrocytic cells did not localize within the nodular compartment. Instead, they were found at the rim of the tumor nodules, forming a transition zone between the two histological MBEN parts (**Fig. 7J,K**). This data suggests that cells that contribute to the tumor microenvironment in MBEN influence different stages of its process.

Taken together, expanded spatial transcriptomic analysis revealed that the tumor microenvironment in MBEN differs between areas of early CGNP-like and neuronally differentiated tumor cells (**Fig. 8**).

**Fig. 8.**
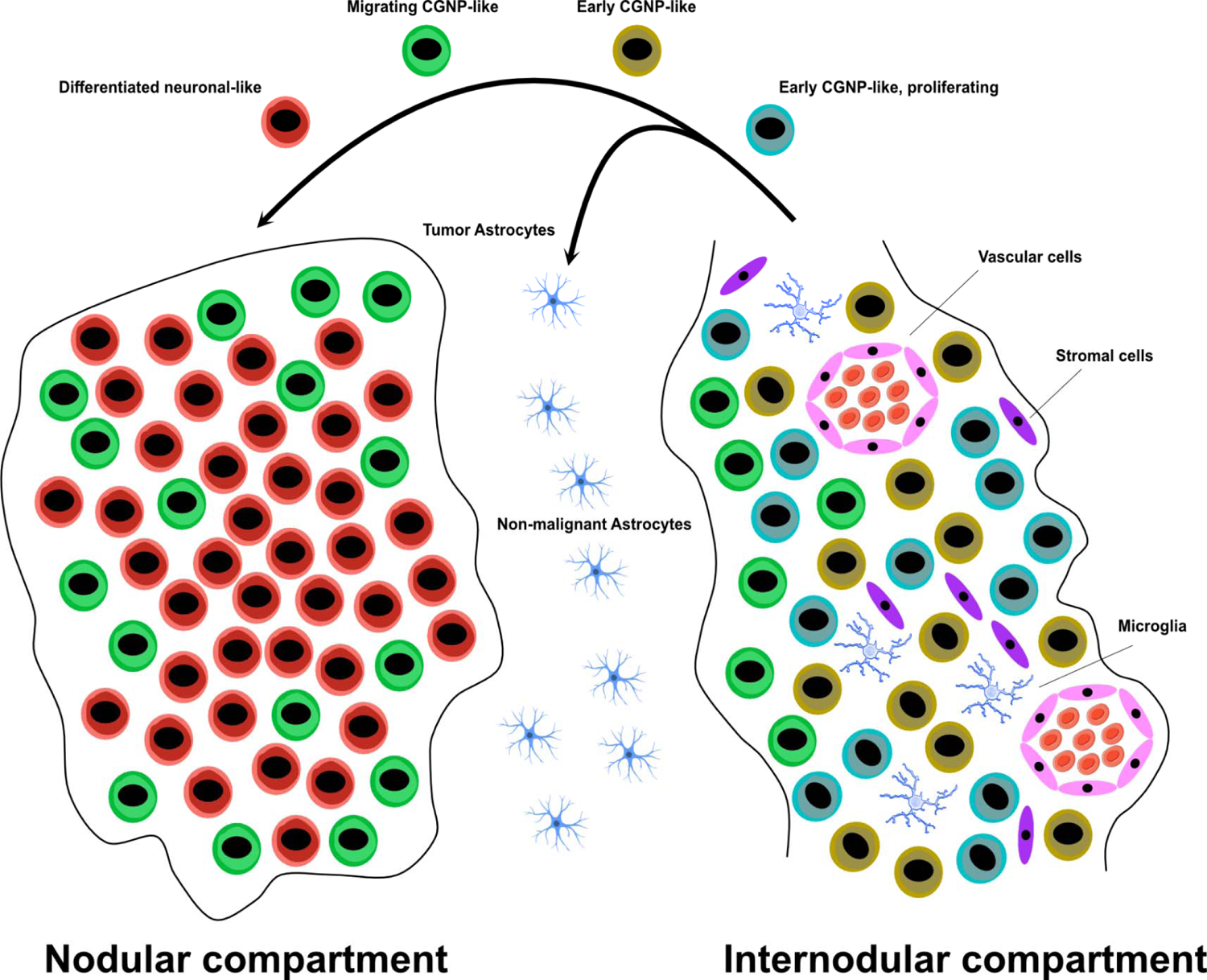
The developmental and spatial architecture of MBEN.

## Discussion

In this study, we applied an integrated multi-modal transcriptomics approach to unravel the underlying biology of the histomolecular heterogeneity of MBEN. Our data indicate that histologically apparent nodular and internodular areas reflect a spectrum of cell stages that are connected through an underlying developmental trajectory which mimics the physiological differentiation of the CGNP-lineage during cerebellar development. Throughout this process, MBEN cells lose proliferative activity and differentiate into a neuronal-like phenotype, which is thought to explain favorable prognosis in most patients including case reports of maturation into benign gangliocytomas.^23, 24^ Our results are supported by an accompanying study from Gold *et al.*, who identified the same MBEN lineage in an independent snRNA-seq dataset and used spatial proteomics assays to observe similar patterns of cellular organization as in our spatial transcriptomics experiments. These insights are of high interest given that a distinct biological property of pediatric cancer is a block of developmental maturation rather than gained ability to de-differentiate. In line with this conclusion, (epi-)genetic maturation blocks have been identified as an emerging topic in pediatric oncology representing potential targets for new therapeutic avenues.^70^ Our data suggest that reactivation of CGNP-associated signaling pathways mimicking normal development and distinct intercellular communication processes result in this phenomenon. It remains enigmatic though, why these processes differ in more aggressive variants of SHH-MB, such as *TP53-*mutated SHH-MB. Interestingly, Gold *et al.* find evidence that late stage, neuronally differentiated MB cells are more common in MBEN than in more aggressive SHH-MB, indicating a potential maturation block in the developmental lineage of these cells. One hypothesis is that tumor formation in MBEN largely depends on SHH-pathway activation in proliferating CGNPs, but without the potential to form a more aggressive phenotype.^15^ The consistent upregulation of SHH-mediated transcription factors may be strong enough to induce neoplastic transformation in early CGNPs in the EGL, which exhibit high proliferative and migratory potential even under physiological circumstances. Similar to early CGNPs, undifferentiated MBEN cells exhibited upregulation of SHH-signaling, whereas late stage MBEN cells showed activation of cellular processes such as Ca^2+^- and NMDA receptor-signaling, which are characteristic for more differentiated CGNPs in the developing cerebellum.^71^ These findings indicate that shared molecular functions between embryogenesis and MBEN formation may exist. Furthermore, it seems likely that epigenetic regulation, for instance via the PRC2-complex, may play a role in MBEN-differentiation, which is underlined by the fact that *EZH2* was upregulated in proliferating, early CGNP-like MBEN cells.^42, 45^

Several studies recently applied single cell RNA-sequencing to MB.^27, 28, 39, 72^ Hovestadt *et al.* reported that infant SHH-MB showed transcriptional similarity to intermediate and late stage CGNPs, while adult tumors correlated with undifferentiated CGNPs and unipolar brush cell-progenitors.^27^ Interestingly, studies by Hovestadt *et al.* and Vladoiu *et al.*, which focused mainly on group 3/4-MB, did not report on the existence of tumor cells with astroglia signatures, while this cell type was also identified in a study on murine SHH-MB models by Ocasio *et al.*.^39^ This raises the intriguing hypothesis that tumor astrocytes associated with MB may be characteristic for SHH-MB. One histopathologic study reported that astrocytic differentiation and GFAP-positivity were restricted to MB cells from nodular tumors and not present in classic MB, however, this study dates back to a time when MBEN was not yet established as a distinct entity.^73^ The hypothesis of CGNP-derived tumor cells developing into tumor astrocytes is supported by the fact that CGNPs can differentiate into astroglial cells upon exposure to elevated levels of SHH.^74^ The original publication that coined the term “MBEN” by Giangaspero *et al.* already described astrocytic, GFAP-positive tumor cells in the internodular compartment, which is why we suggest giving MBEN-associated tumor astrocytes the byname “Giangaspero cells”.^16^

Microglia, astrocytes, and astrocyte-like tumor cells play an important role in the formation and progression of SHH-MB.^38, 40, 75–77^ We attempted to distinguish between malignant and non-malignant astrocytes based on their clustering behavior and CNVs. This approach was without alternatives in light of the current lack of MBEN models, and future investigations will be needed to further unravel the functional and genomic differences between these two cell types in the human setting. However, several studies have generated convincing evidence that both normal and tumor-derived astrocytes may support SHH-MB proliferation and growth in murine models.^38, 75–77^ Our data indicate that cells with an astrocytic phenotype are found at the border between the two compartments, co-localizing with MBEN cells in their migrating and later, postmitotic differentiation stages. In contrast, all other non-malignant cell types were in close proximity to early CGNP-like tumor cells. Given the fact that astrocytes are known to influence surrounding cells via paracrine signaling, these patterns suggest that astrocytic cells may have a direct influence on the differentiation process of MBEN, whereas other non-malignant cell types are mostly involved in supporting its proliferating subpopulations, or, in the case of microglia, trying to suppress tumor growth.^78^ With these observations in mind, the internodular compartment of MBEN can be understood as a source of neoplastic cells, which then migrate into the nodular compartment while losing proliferative potential.

DNMB, another histological variant of SHH-MB, also shows a bicompartmental structure, but represents a biologically and clinically distinct phenotype. Results of Gold *et al.* indicate that the nodular areas may vary in their level of differentiation, with MBEN tumors being more likely to contain cells resembling later stages of differentiation. To date, murine or cell models that faithfully preserve the unique histological composition of MBEN and could be used to perform functional analyses are lacking.

Our study resolves the intratumoral heterogeneity of MBEN at the single cell level and reveals the spatial relation between the different cell types in the context of its bicompartmental histological structure. Thus, it provides a framework for similar analyses in other malignancies with intratumoral heterogeneity.^79^ Finally, further deepening our understanding of the biological principles underlying both intratumoral differentiation processes and maturation blocks are expected to guide the development of drugs that either induce or overcome these phenomena in embryonal cancer types.

## Methods

### Material and data collection

All experiments in this study involving human tissue or data were conducted in accordance with the Declaration of Helsinki. Cases from earlier studies on MBEN were screened regarding the availability of fresh frozen tissue.^15, 25, 80^ Clinical data and tissue of all nine cases in the study were collected from patients from the international DKFZ cohort after receiving written informed consent from the respective patients or their legal representatives and after approval by the ethic boards of Heidelberg University and the German Cancer Research Center.

### DNA-methylation profiling and CNV analysis

DNA methylation profiling was performed using the Infinium Human Methylation 450k and EPIC BeadChips as previously described.^9^ Subsequently, the Heidelberg Brain Tumor Methylation Classifier v11b6 (https://www.molecularneuropathology.org) was applied for molecular classification. Copy-number variation analysis from 450k and EPIC methylation array data was performed using the conumee Bioconductor package version 1.12.0 (Hovestadt V, Zapatka M, 2017).

### RNA-bulk sequencing

Bulk RNA-seq data were analyzed as described previously.^80^ Shortly, reads alignment was performed with STAR tool^81^ and gene expression counts were computed with the Subread package.^82^

### DNA-panel sequencing

Molecular barcode-indexed ligation-based sequencing libraries were constructed using 200 ng of sheared DNA. Libraries were enriched by hybrid capture with custom biotinylated RNA oligo pools covering exons of 130 cancer-associated genes. Paired-end sequencing was performed using the NextSeq 500 (Illumina). Sequence data were mapped to the reference human genome using the Burrows–Wheeler Aligner and were processed using the publicly available SAM tools. Only variants annotated as “exonic” or “splicing” were included, “intergenic” and other untranslated regions were excluded. Recurrent gene mutations of PTCH1, SUFU, and SMO were also assessed with residual DNA from the same pool used for sequencing by polymerase chain reaction followed by direct Sanger sequencing of the corresponding exons.

### Nuclei isolation

Prior to nuclei isolation^83^, all necessary materials (douncer and pestles, falcon tubes, pipette tips) were pre-coated using coating buffer (sterile phosphate buffered saline (PBS), filtered with Millex HA filter units 0.45 µl). Fresh frozen tissue was placed on dry ice and cut with a scalpel. Next, the tissue was put into 5ml of lysis buffer containing DTT and Triton-X (0.1 %), and subsequently lysed using a glass douncer and pestle. The nuclei were filtered two times (100µM and 40µM filters) and centrifuged for 5min at 1600rpm and 4°C, followed by two washing steps (5min, 1600rpm, 4°C) using a PBS-containing washing buffer. The resulting pellet was resuspended in 1ml of storage buffer. Nuclei were checked for integrity and the absence of cell debris under a light microscope and subsequently counted using a Luna Automated Cell Counter (Logos Biosystems).

### SnRNA-seq 10X Genomics

The 10X Genomics 3’-Single Cell RNA-sequencing V2 protocol was applied to all samples according to the manufacturer’s instructions using 10X Chromium Single Cell 3’ Reagent Kits v2 (https://www.10xgenomics.com/). 14,000 nuclei were loaded per sample and processed with the Chromium Controller. The resulting cDNA-libraries were quantified with Qubit fluorometric quantification (ThermoFisher Scientific) and quality assessment was done using the TapeStation system (Agilent). Sequencing was performed according to the manufacturer’s instructions.

### SnRNA-seq SMARTseq V2.5

Whole transcriptome snRNA-seq was performed following an adapted SMARTseq protocol (SMARTseq V2.5) for nuclei.^84^ For each sample, single nuclei were sorted in 384-well plates into 1 µl of lysis buffer containing RNaseinhibitors and a polyT oligo. For the reserve transcription reaction, we used a TSO with LNAs (Eurogentec) at a final concentration of 3 µM and the Maxima H Minus reverse transcriptase (Thermo) under molecular crowding conditions with 7.5% PEG-8000. Subsequently, PCR was performed with the KAPA HiFi HotStart ReadyMix (KAPA Biosystems), followed by a purification step with AMPure XP beads (Beckman) using the Agilent Bravo system. Library preparation was done via the Nextera XP Library Prep kit (Illumina) using 350 ng of cDNA as input. The reaction volume of each step was reduced by a factor of three and all pipetting steps were done with the mosquito LV (SPTlabtech). After pooling all libraries from one plate, quantification and quality control using the Qubit fluorometric quantification (ThermoFisher Scientific) and TapeStation system (Agilent) was performed, and the cDNA libraries were sequenced on a Nextseq550.

### Analysis of snRNA-seq raw data

Initial processing of 10X data (reads alignment, counts computation per cell) was performed with CellRanger v3 pipeline. The SMARTSeq V 2.3 (SS2.5) data was processed per sample representing a cell with bulk RNA-seq derived procedure as described previously.^80^ For both protocols hg19 human genome reference combined with gencode v19 gene annotation were used. For gene expression counts computations exons were merged with introns due to single nuclei protocol specificity.^85^

Computed cell gene expression counts were further analyzed per sample with Seurat toolkit.^86^ Filtering control limits (minimum number of molecules/genes) were identified based on the quality control inspection. The doublets detection was performed with the DecontX tool.^87^ Copy number profiling was performed with InferCNV.^36, 88^ Signatures for Ca^2+^ and BMP signaling were constructed using the module score function within Seurat. Gene ontology analysis was performed using the PANTHER classification system with DEGs per cluster as input (http://www.pantherdb.org/).^61^ The results were visualized with the REVIGO package by clustering based on semantic similarities.^89^

Initially 10X and SS2.5 datasets were analyzed separately; their identified clusters were merged into pseudobulk blocks and compared via correlation. Non-tumor cells were further annotated based on inspection of clusters from merged datasets: the clusters containing cells from multiple samples were considered as normal while consisting from one sample on 95% as a tumor. Afterwards, the datasets (full and tumor only) were merged using Harmony package^31^ considering both sample and protocol as batch effects. Trajectories were identified from application of Slingshot R package.^90^ The similarity of tumor cells to normal cerebellum cell types was analyzed using the SingleR package.^32^

For Transcription factor (TF) activity estimation the python version of DecoupleR package was used.^91^ The needed conversion from Seurat objects to AnnData objects was done with the SeuratDisk library (https://mojaveazure.github.io/seurat-disk/). TF activity estimation was performed based on DoRothEA which is a comprehensive prior knowledge resource containing curated TFs and their targets.^64^ This network was derived from the OmniPath database^92^ via DecoupleR. In DoRothEA, each TF-target interaction includes a confidence level annotation ranging from A to E based on the supporting evidence where A is the highest confidence level and E is the lowest. For this analysis, TF-target pairs coming from the three highest confidence levels (i.e., A, B and C) were used to create a predictive model for TF activity estimation. The estimations were done with a multivariate linear model based on the log transformed gene expression data. The resulting activities were summarized per cluster by their mean with the summarize_acts function. The minimum standard deviation was set to zero to retrieve all results.

### Microdissection

Microdissection with subsequent RNA isolation was performed on FFPE-derived histological slides from 26 MBEN patients as previously described.^93^ The main analysis of microdissected samples was performed as for standard bulk RNA-sequencing.^80^ The clustering was performed from normalized gene expression counts with a focus on top 500 highly variable genes. Differentially expressed genes between nodular and inter-nodular blocks were detected via limma package, with considering the tumor sample for batch effect correction. Further, nodular and inter-nodular detected groups of differentially expressed genes were used as a reference for Gene Set Variation Analysis^69^ on the genes identified as markers of clusters in MBEN single cell data.

### smRNA-FISH (RNAScope)

#### Histological sample preparation

Sectioning of fresh/frozen tissue derived from patients MB266, MB295 and MB299 was performed at −20°C on a cryostat (Leica) and 8-10µm sections were mounted on Superfrost Plus slides (ThermoFisher). Cryosections were stored at −80°C until further use.

#### Virtual H&E staining and probe hybridization

12-plex single molecule RNA-FISH (smRNA-FISH) was performed using the RNAScope HiPlex assay (ACDbio/biotechne) as described in the ‘RNAScope HiPlex Assay User Manual (324100-UM)’ with minor adaptions (**Suppl. Tbl. 7**). Briefly, sections were fixed in 4% paraformaldehyde (PFA) for 60 min, washed two times with PBS and dehydrated in Ethanol. For virtual H&E staining (PMID: 29531846), sections were stained with Eosin (Sigma, 1:10 diluted in 0.45 M Tris acetic acid, pH=6) for 1 min at room temperature, washed in H_2_O and incubated for 15 min in 4x SSC buffer. Sections were then stained with DAPI for 30 sec and mounted in Prolong Gold Antifade (ThermoFisher). After the first virtual H&E imaging round (R0), the coverslip was removed by incubation in 4x SSC buffer for 15-30 min. Afterwards, sections were washed in PBS once and again dehydrated in Ethanol. Sections were then treated with Protease IV (ACDbio) for 30 min at RT, washed 2x with PBS and incubated with transcript-specific (Table 1) and amplifier probes according to the manufactureŕs instructions. Between imaging rounds, fluorophores of the previous imaging rounds were cleaved to enable consecutive rounds of imaging, with each round targeting a new set of transcripts. Up to four transcripts were labeled per imaging round by Alexa488, Atto550, Atto647 and Alexa750 fluorescent dyes. For MB266 and MB299, four transcripts (+DAPI) were imaged in three imaging rounds (R1-R3). For MB295, three transcripts (Alexa488, Atto550, Atto647 and DAPI) were imaged in four imaging rounds using the ‘RNAScope HiPlex Alternate Display Module’ (R1-R4).

#### Microscopy

smRNA-FISH images were acquired on an Andor Dragonfly confocal spinning-disk microscope equipped with a CFI P-Fluor 40X/1.30 Oil objective. The region of interest was selected based on the DAPI signal and 50 z-slices were acquired with a step size of 0.4 µm (20 µm z-range) per field of view (FOV). Lasers and filters were set to match fluorescent properties of DAPI and above mentioned dyes. Tiles were imaged with a 10% overlap to ensure accurate stitching.

#### Image analysis

Pre-processing of images was performed in ImageJ. Images were projected in z using Maximum Intensity Projection. Illumination correction was performed on the DAPI and Eosin images for virtual H&E visualization using the BaSiC plugin.^94^ Image tiles were stitched using ‘Grid/Collection stitching’ and registered afterwards based on the DAPI signal using ‘Register Virtual Stack Slices’ using the Affine feature extraction model and the Elastic bUnwarpJ splines registration model. Virtual H&E transformation of DAPI and Eosin staining was performed in R using the EBImage tool and custom script.^95^ Nuclei were segmented by the Cellpose python tool using the ‘cyto’ model and a model match threshold of 1.5.^96^ Nuclei outlines were exported to ImageJ and transformed to ROIs using the ROImap function of the LOCI plugin. Spot detection was performed using the RS-FISH plugin in ImageJ (https://doi.org/10.1101/2021.03.09.434205) with Sigma set to 0.93. The threshold for spot detection was adapted for each patient and fluorophore individually (range: 0.004 – 0.0086). Transcripts were assigned to nuclei using a custom KNIME script that overlaps the DAPI and spot signal and counts spots per nuclei. Nuclei metadata including x/y coordinates and other features were extracted using the ‘Segment Features’ node in KNIME.

#### Downstream analysis

Raw transcript count matrices were generated from KNIME outputs in R using custom script. Transcript counts for *LRRTM4* and *FOS* were removed from the analysis due to strong discrepancies to snRNA-seq data. smRNA-FISH data was further analyzed with the Seurat tool^97^: Cells were filtered by transcript count (<5 and >100 transcripts) as well as by nuclei size (<90 and > 2000 pixels). smRNA-FISH data was normalized using scTransform^98^, visualized using principal component analysis and clustered using the Louvain algorithm. Clusters with similar marker gene expression and high correlation of their average gene expression profiles were merged and assigned to cell types according to their marker expression as detected in snRNA-seq data. Further spatial analysis and visualization of smRNA-FISH data including spatial networks was performed using the Giotto tool in R.^99^ Cell type-specific nuclei mask images were generated using KNIME and visualized with napari (https://napari.org).

### Molecular Cartography (Resolve Biosciences)

To ensure that both spatial transcriptomic methods were comparable, smRNA-FISH marker genes were included in our MB panel, alongside marker genes of the developing cerebellum, oncogenes, and markers of non-malignant cell types. OCT-embedded samples were cryo-sectioned as described above into 10 µm sections onto an MC slide. Fixation, permeabilization, hybridization and automated fluorescence microscopy imaging were performed according to the manufacturer’s protocols (Molecular preparation of human brain (beta), Molecular coloring, workflow setup) except for a few adaptations indicated below. Briefly, slides were thawed at room temperature and dried at 37°C. Subsequently, the MC observation chamber was assembled by attaching sticky wells (8-well) to the MC slide. Sections were fixed, permeabilized, rehydrated and treated with TrueBlack (Biotium) autofluorescence quencher. In contrast to the MC protocol provided by Resolve Biosciences, we diluted the quencher 1:20 as specified by Biotium. Next, the sections were thoroughly washed and primed before the specific probes against our genes of interest were hybridized at 37°C overnight. The probe sequences were designed by Resolve Biosciences’ proprietary algorithm and are hence not listed here. After hybridization, the sections were washed, and the MC observation chamber was transferred to the MC machine for eight automated iterations of coloring and imaging to decipher the transcript localization of the 100 different genes of interest in the tissue.^100^ Therefore, regions of interest (ROIs) were selected for each section (MB263, MB266, MB295, MB299) based on a brightfield overview scan. In the last imaging round, Nuclei were stained with DAPI yielding a reference image for nuclei segmentation. After the run, the MC software performs registration of the raw images, assigns transcripts to the combinatorial color codes detected and combines individual tiles to ROI panoramas. The outputs are text files containing the transcript coordinates in 3D as well as maximum projections of DAPI images for each ROI.

After the MC run, we stained the tissues with an anti-NCAM antibody and eosin for virtual H&E images as described above. All following steps are carried out at room temperature and washing was always done 3x with PBS unless specified otherwise. Sections were washed, fixed in 4% paraformaldehyde (PFA) for 30 min, washed again, permeabilized with 0.2% Triton X-100/PBS for 12 min and washed again. Next, the sections were blocked for 1 h in 10% goat serum/PBS and then incubated with a 1:200 dilution of the primary antibody (mouse anti-NCAM/CD56 from ThermoFisher Scientific, MA1-06801, lot: 17425) in 10% goat serum/PBS for 1 h. After washing 3x 5 min with 0.002% NP-40/PBS, the sections were incubated with a 1:300 dilution of the secondary antibody (goat anti-mouse Alexa568) in 10% goat serum and washed again before and after a 15 min incubation with 5 µM DAPI. After washing 1x with double-distilled water, the sections were incubated with Eosin mix (1 vol Eosin Y Sigma HT110216 to 9 vol Tris Acetic acid buffer 0.45 M, ph6) for 1 min. After ten washes with double-distilled water, MC imaging buffer was added to the sections and they were imaged on the Andor Dragonfly microscope described above with the following setup: 60x objective, 41 *z*-slices at 0.3 µm distance, 10% overlap between tiles for stitching, EM gain 200, excitation at 405 nm (DAPI), 488 nm (Eosin) and 647 nm (immunofluorescence) with appropriate emission filters.

DAPI images were flatfield-corrected using a separately recorded flatfield and darkfield image. Stitching and registration of the Dragonfly DAPI image to the MC DAPI image (from round 8) and the calculation of the virtual H&E images was done as described above.

The detection of cell boundaries was performed with QuPath.^101^ Afterwards, gene expression counts were computed per cell and extracted using Resolve Bioscience plugin in the ImageJ2 toolkit.

Formed gene expression matrix analysis (filtering, clustering, visualization) was performed with Seurat toolkit.^86^ The samples were merged using the Harmony R package with batch effect adjustment.^31^ Additional spatial analysis (closest cell connections detection) was performed with Giotto toolkit analogous to the analysis of the smRNA-FISH dataset.^99^

### Statistical analysis and visualisations

The Kaplan-Meier-method was applied to analyze and visualize progression free and overall survival. The respective analysis were performed using the R packages survival (https://github.com/therneau/survival) and survminer (https://github.com/kassambara/survminer). Descriptive statistics and visualizations were conducted using the R base and ggplot2 packages.

## Acknowledgements

We thank the High-Throughput Sequencing Unit of the Genomics and Proteomics Core Facility, German Cancer Research Center (DKFZ) for providing excellent services regarding all sequencing experiments. This project received funding by a grant of the European Research Council (ERC) to S.M.P. under the European Union’s Horizon 2020 research and innovation program as part of the initiative BRAIN-MATCH (Grant agreement ID: 819894). D.R.G. was supported with personal grants by the German Academic Scholarship Foundation (Studienstiftung des Deutschen Volkes) and the Mildred Scheel Doctoral Fellowship program of the German Cancer Aid (Deutsche Krebshilfe). A.S.R. and J.S.-R. were supported by the German Federal Ministry of Education and Research (BMBF) through the grant CompLS DeepSC2 (031L0269B). D.R.G. and K.W.P. are thankful to the non-profit foundation Ein Kiwi gegen Krebs for their support. In order to design Fig. 4 and Fig. 8, the two images “Blue Astrocyte” (Andrew Hardaway) and “Microglia Resting” (John Chilton) from the database https://scidraw.io/ were used.

## Author contributions & Competing Interests

Acquisition, analysis, and interpretation of data: D.R.G., K.O., A.R., S.T., K.K.M., J.S., B.S., A.S.R., K.B., S.S., M.B., F.G., K.E., J.S.-R., D.T.W.J., D.K., J.-P.M., K.R., A.K., S.M.P., K.W.P. Design and conceptualization: D.R.G., K.O., A.K., S.M.P., K.W.P. Writing of the original draft: D.R.G., K.O., A.K., S.M.P., K.W.P. Substantial revisions and feedback to the draft: All authors. S.T. is an employee of the company Resolve BioSciences GmbH. J.S.-R. has received funding from GSK and Sanofi as well as consultant fees from Travere Therapeutics and Astex Pharmaceuticals. No other author declares a conflict of interest.

## Supplementary Figures

**Suppl. Fig. 1.**
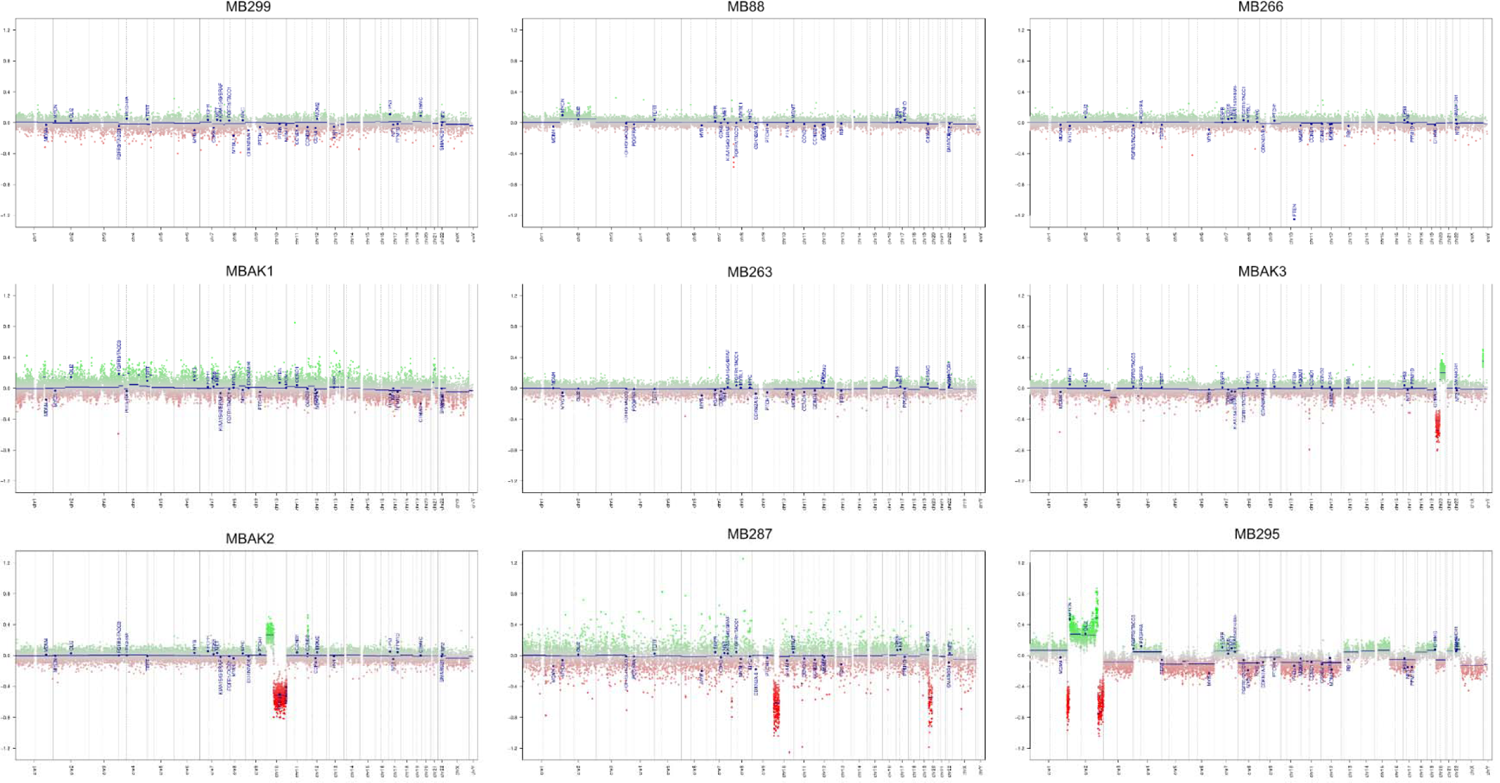
CNVs are infrequent in MBEN. CNV-plots derived from DNA-methylation profiling data for all nine samples (no chromosomal CNAs: five cases; 1-2 chromosomal CNAs: three cases; >2 chromosomal CNAs: one case)

**Suppl. Fig. 2.**
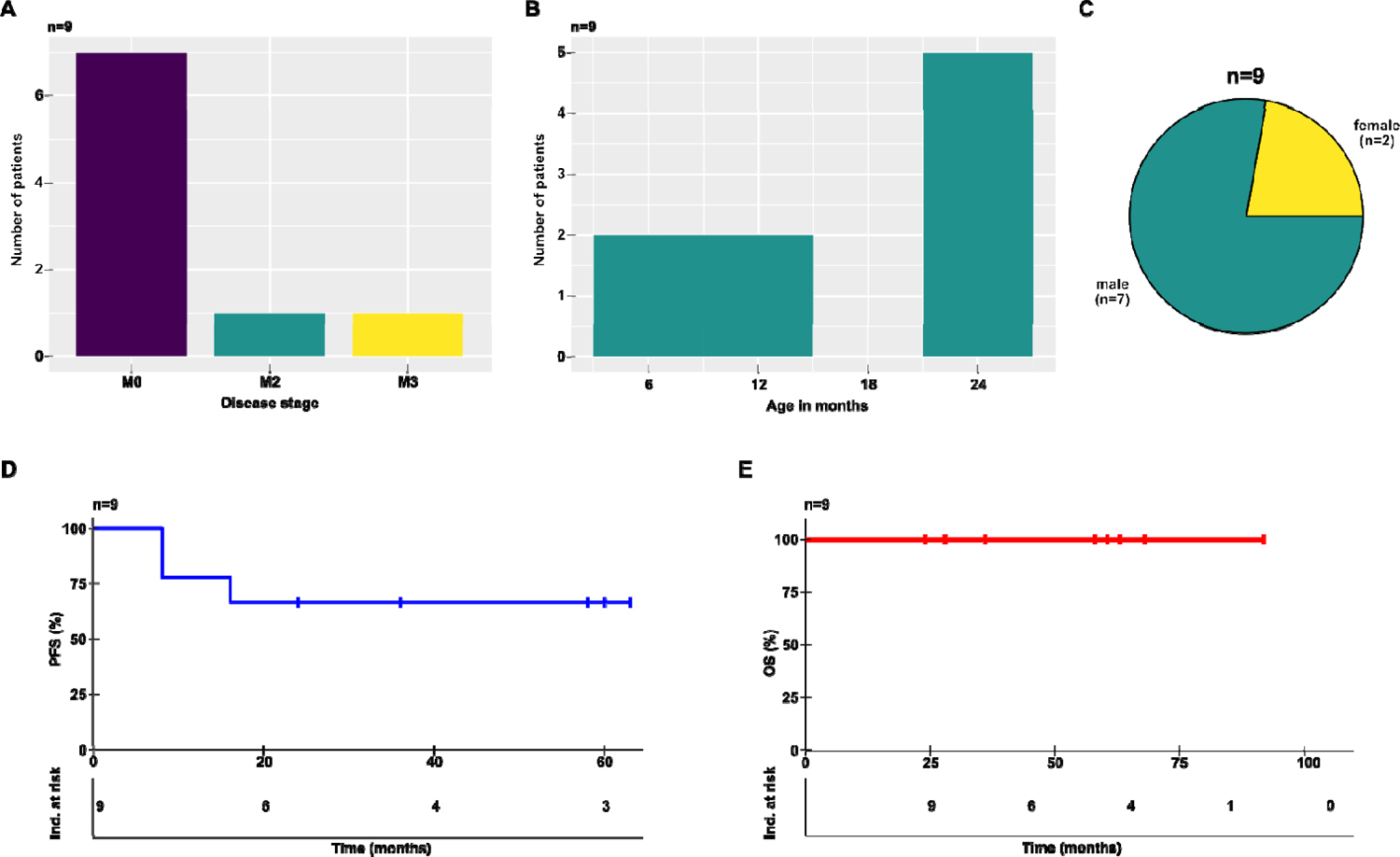
Clinical and epidemiological features of the MBEN patient cohort. **A** Bar chart depicting disease stage at time of diagnosis **B** Histogram of age in years at diagnosis **C** Pie chart of the sex distribution of the cohort **D** Kaplan-Meier-curve showing PFS **E** Kaplan-Meier-curve showing OS. PFS = progression free survival, OS = overall survival

**Suppl. Fig. 3.**
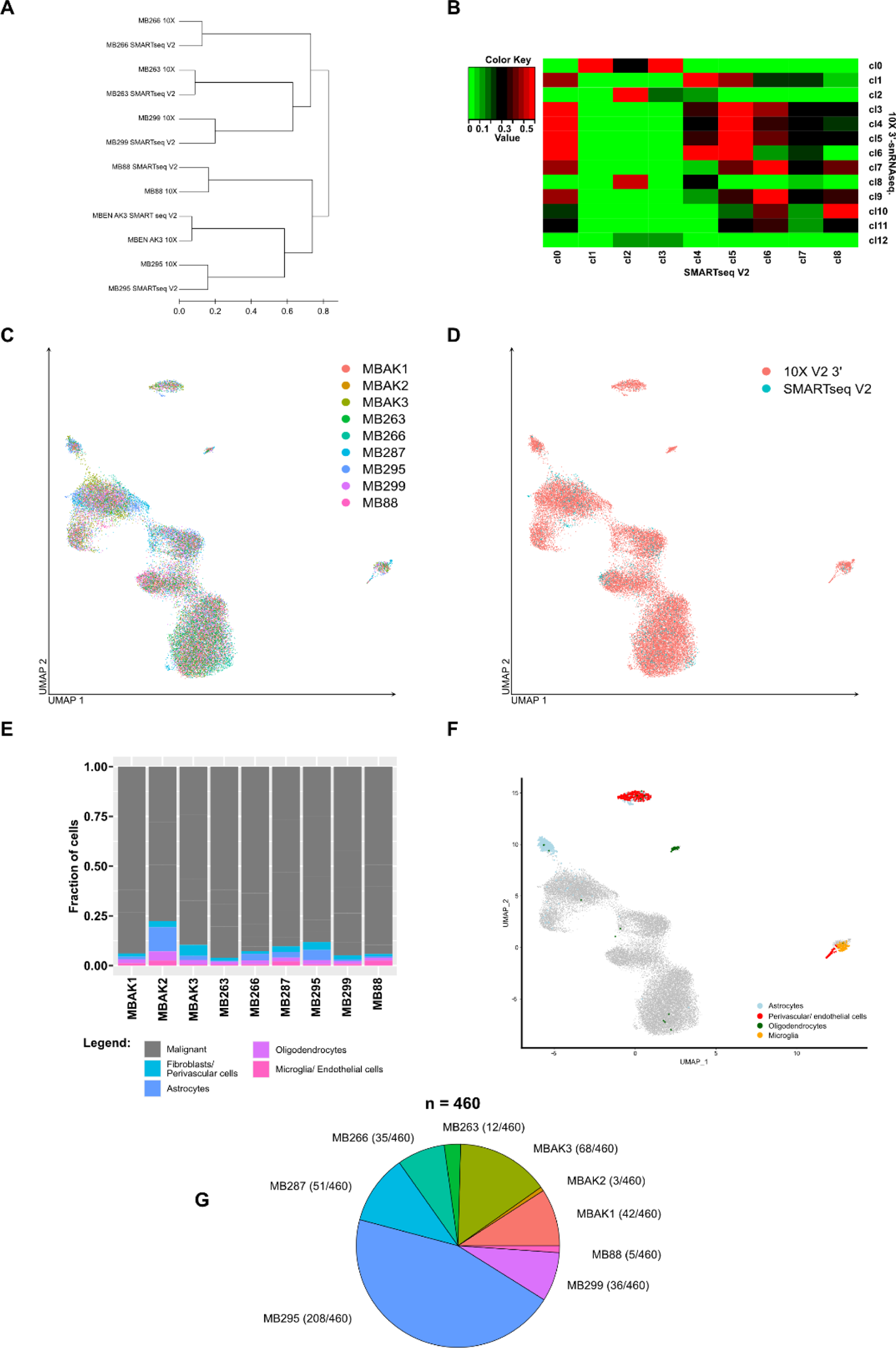
Quality control and identification of non-malignant cell types based on snRNA-seq derived CNV-analysis. **A** Unsupervised clustering of 10X and SMARTseq V2.5 genotype profiles confirms strong transcriptional similarity between clusters derived from both snRNA-seq methods. **B** Heatmap showing clear correlation between clusters of single cells sequenced with two complementary methods (x-axis: SMARTseq. V2.5, y-axis: 10X Genomics). **C** UMAP-projection colorized according to patient. No patient-specific batch effects were detectable after integration. **D** UMAP-projection colorized according to technology confirms good integration across both technological platforms. **E** Stacked bar chart visualizing the distribution of cells across malignant and non-malignant clusters for every patient. **F** Non-malignant cells were mapped onto an atlas of the developing cerebellum to confirm the designation of non-malignant cell types (microglia, astrocytes, oligodendrocytes, perivascular and endothelial). **G** Pie chart summarizing the number of tumor astrocytes per sample. All samples are represented to varying degrees.

**Suppl. Fig. 4.**
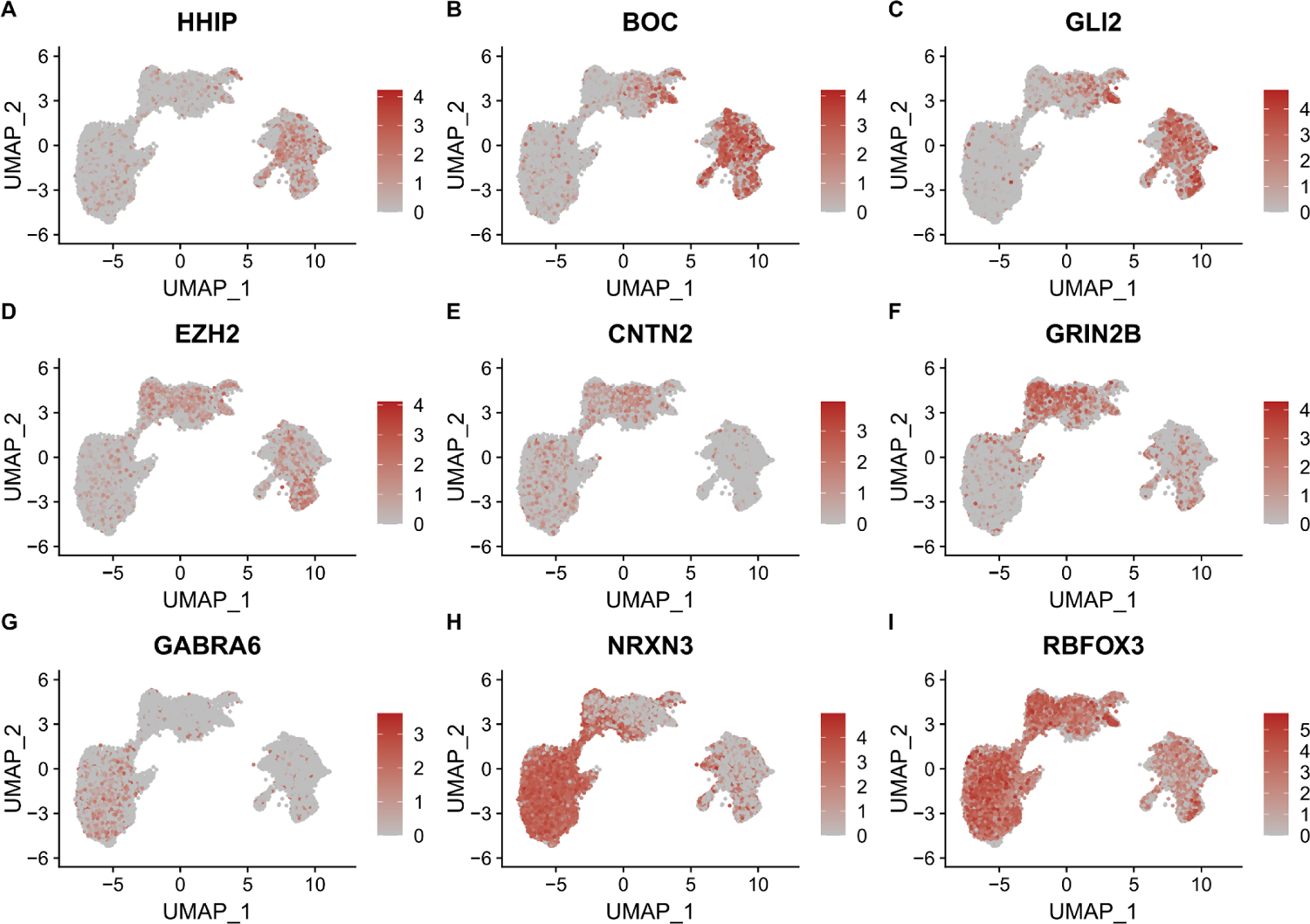
Malignant cell states differ in their expression of CGNP-marker genes. UMAP-plots showing the expression of representative marker genes of CGNP development. **A – C** SHH-pathway members *HHIP, BOC,* and *GLI2*. **D** epigenetic regulator gene *EZH2.* **E** and **F** markers of intermediate CGNP-stages *CNTN2* and *GRIN2B*. **G – I** markers of differentiated CGNPs *GABRA6*, *NRXN3*, and *RBFOX3*.

**Suppl. Fig. 5.**
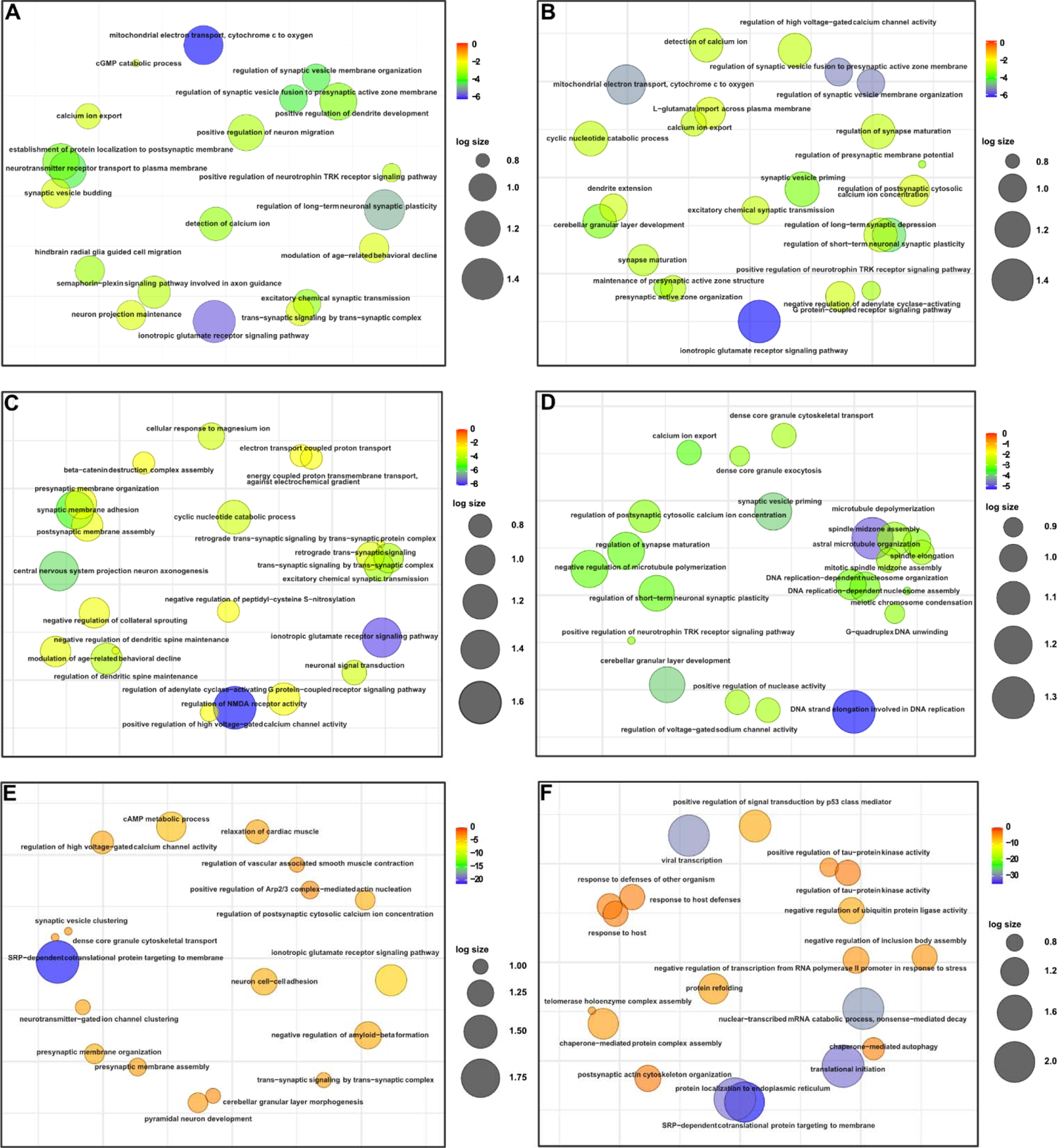
Gene ontology analysis reveals core biological functions of granular cerebellar development to be upregulated in MBEN. Clustering visualizations based on semantic similarity representing the top 25 GO terms based on fold enrichment for **A** Cluster 0, **B** Cluster 1, **C** Cluster 2, **D** Cluster 3, **E** Cluster 4, and **F** Cluster 5. Colorization is based on raw p-values.

**Suppl. Fig. 6.**
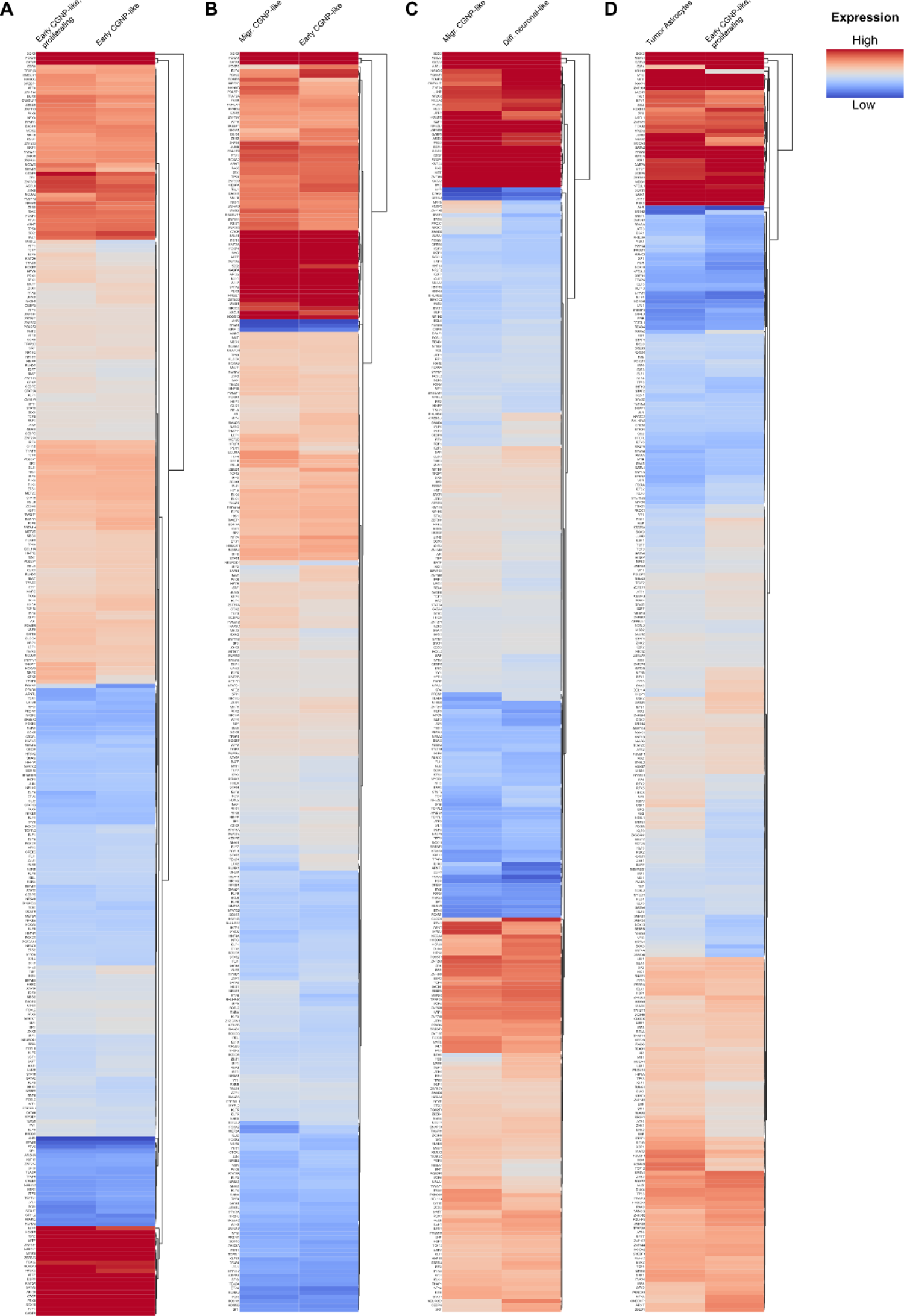
Transcription factor activity changes throughout the course of MBEN development. **A – D** Heatmaps visualizing the changes in TF activity during MBEN differentiation. Each heatmap shows the differential TF activity between two clusters (also compare Fig. 3D and 4E).

**Suppl. Fig. 7.**
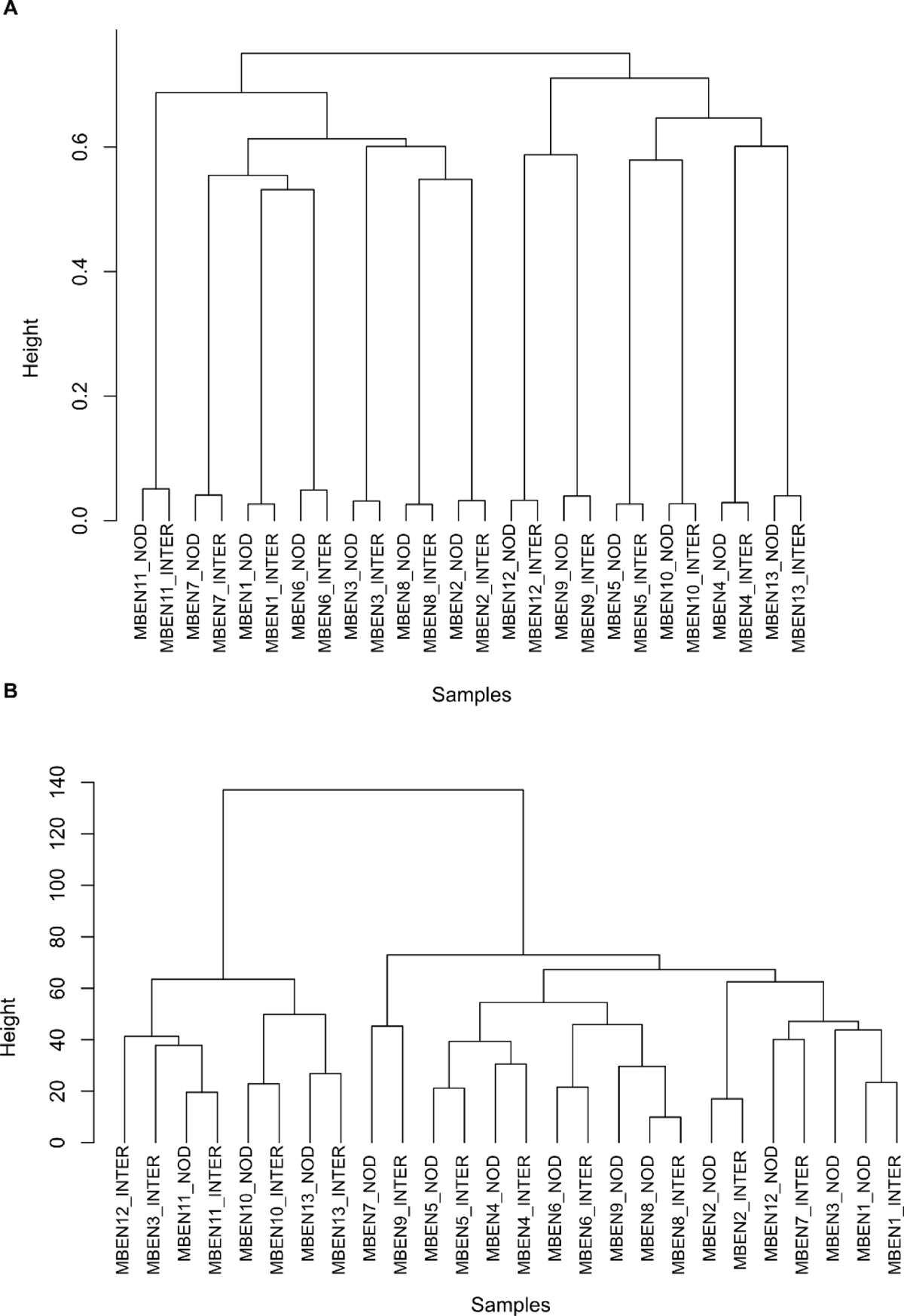
Microdissection datasets quality control and inspection. **A** Unsupervised hierarchical clustering of internodular/nodular profiles using the top 500 most highly variable genes (applied method: ward.D2) **B** Unsupervised clustering of internodular and nodular genotype profiles confirms tissue source from the same tumor sample.

**Suppl. Fig. 8.**
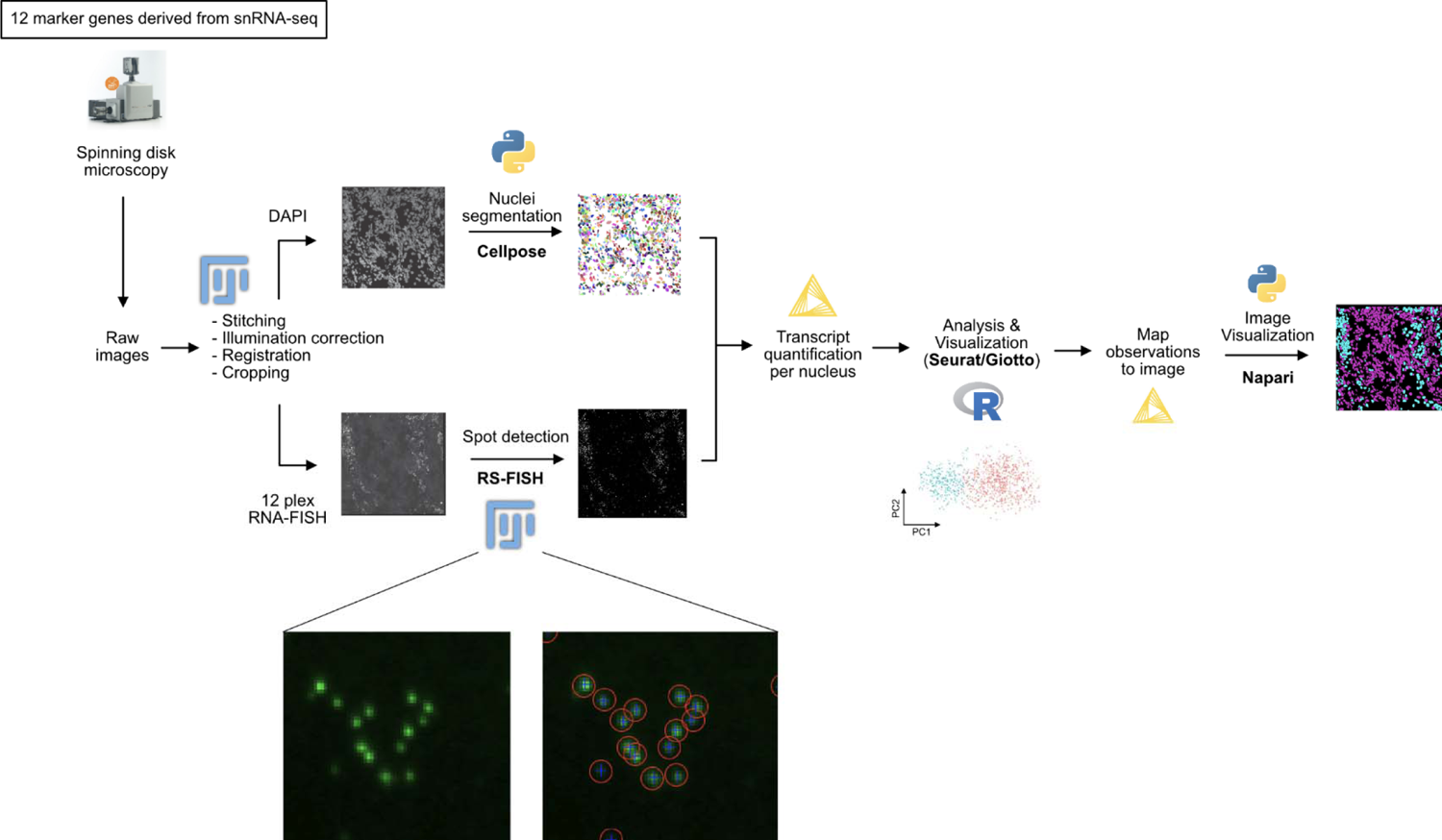
Workflow of the integrated snRNAseq- and RNAscope-analysis. Workflow depicting the single steps of RNAscope analysis and correlation with clusters from snRNA-seq.

**Suppl. Fig. 9.**
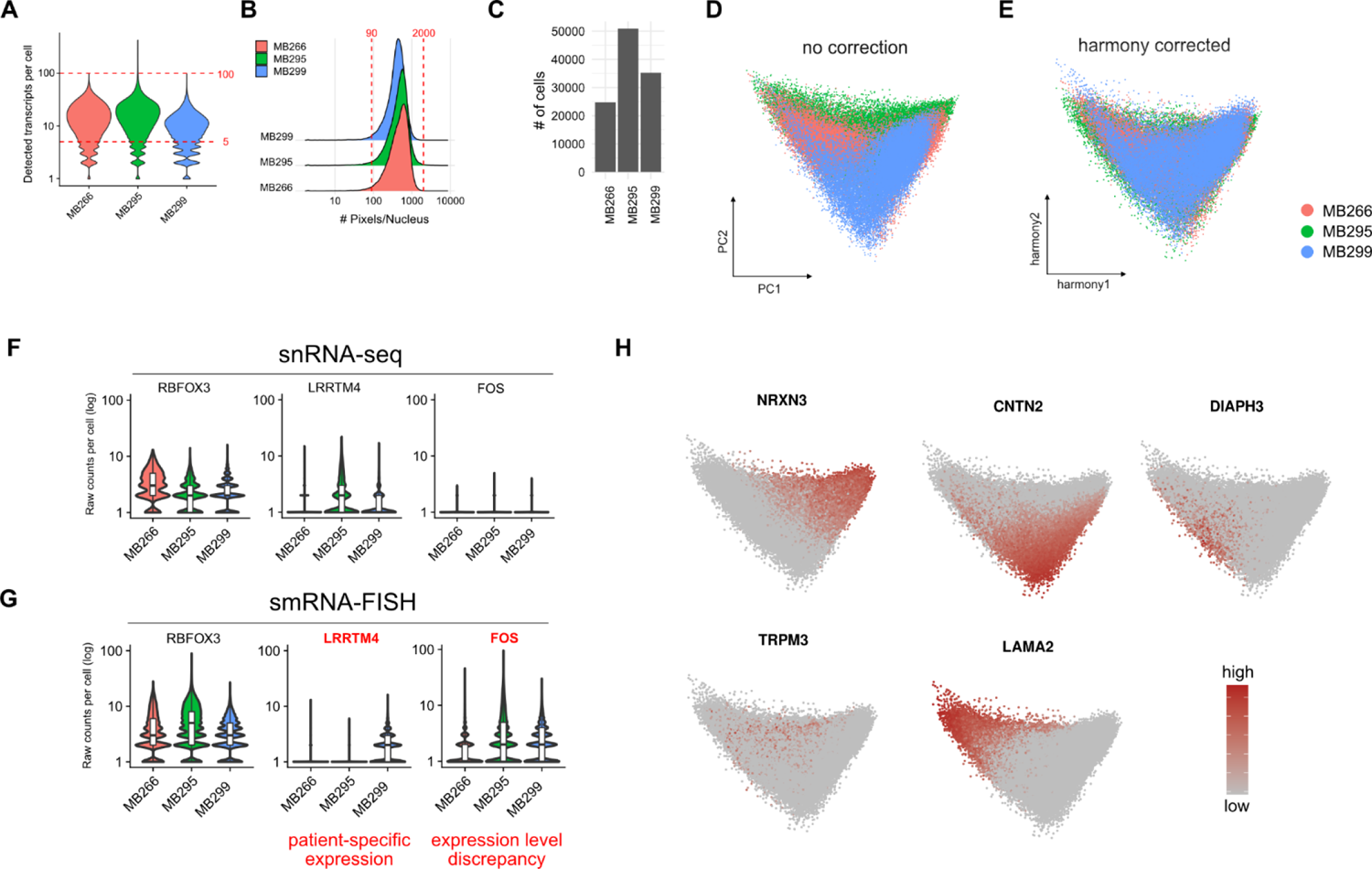
Quality control smRNA-FISH using RNAscope. **A – C** Visualizations showing **A** the number of transcripts per pseudo single cell, **B** the number of pixels per nucleus, and **C** the number of single cells for each sample. **D** Principal component analysis showing high concordance between the three cases even without correction, which is further improved by **E** harmony correction. **F, G** Expression of the three marker genes *LRRTM4, FOS*, and *RBFOX3* in the **F** snRNA-seq and **G** smRNA-FISH datasets, respectively. *LRRTM4* and *FOS* were excluded from the further analysis due to strong expression level discrepancies or patient-specific expression (*RBFOX3* shown for comparison). **H** Expression of snRNA-seq derived marker genes in the smRNA-FISH dataset.

**Suppl. Fig. 10.**
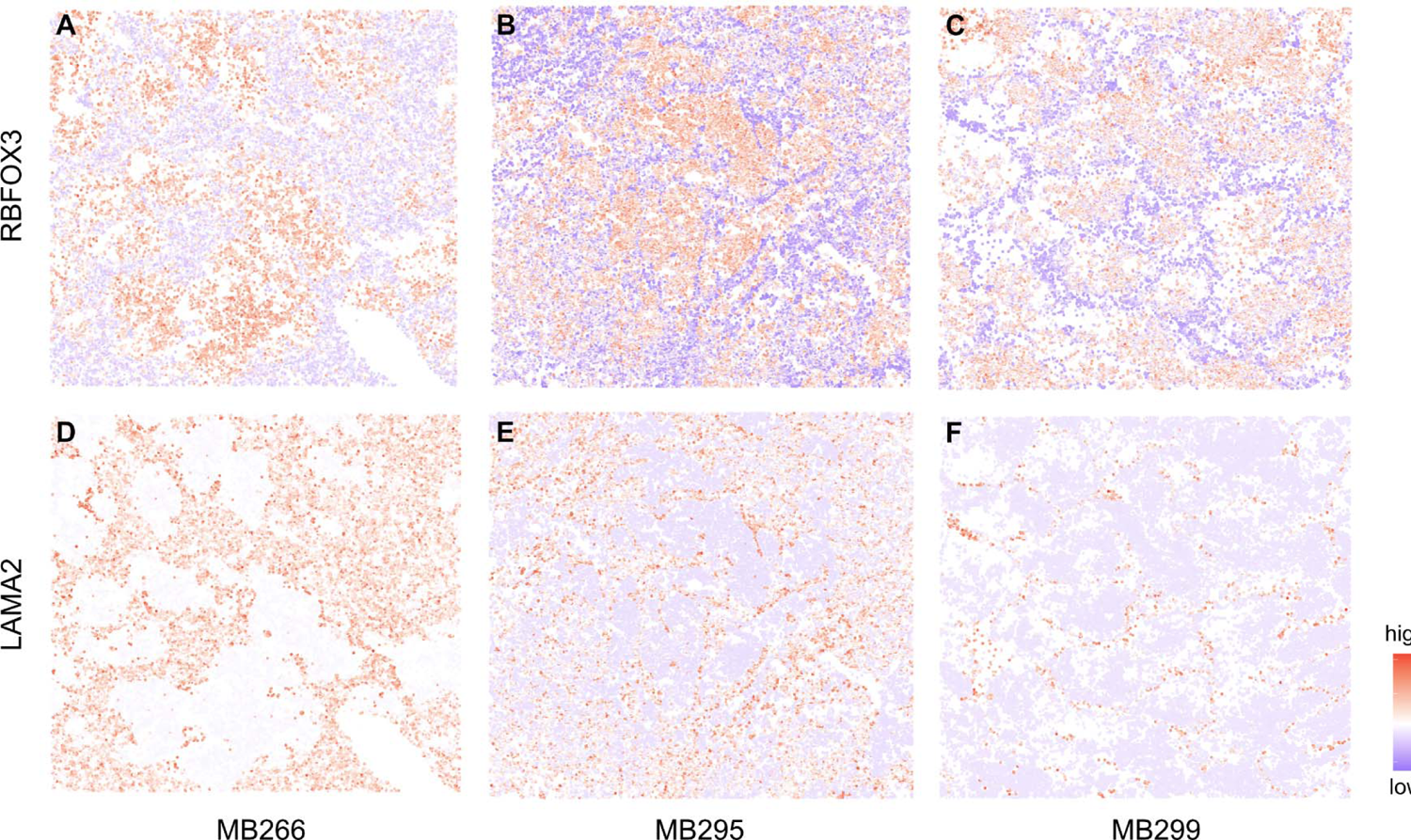
Population-specific marker genes recapitulate MBEN histology. The marker genes *RBFOX3* and LAMA2 are differentially expressed in all three MBEN samples. **A – C** *RBFOX3* is highly expressed in the nodular compartments, while *LAMA2-*expression is restricted to the internodular parts of the respective tumors.

**Suppl. Fig. 11.**
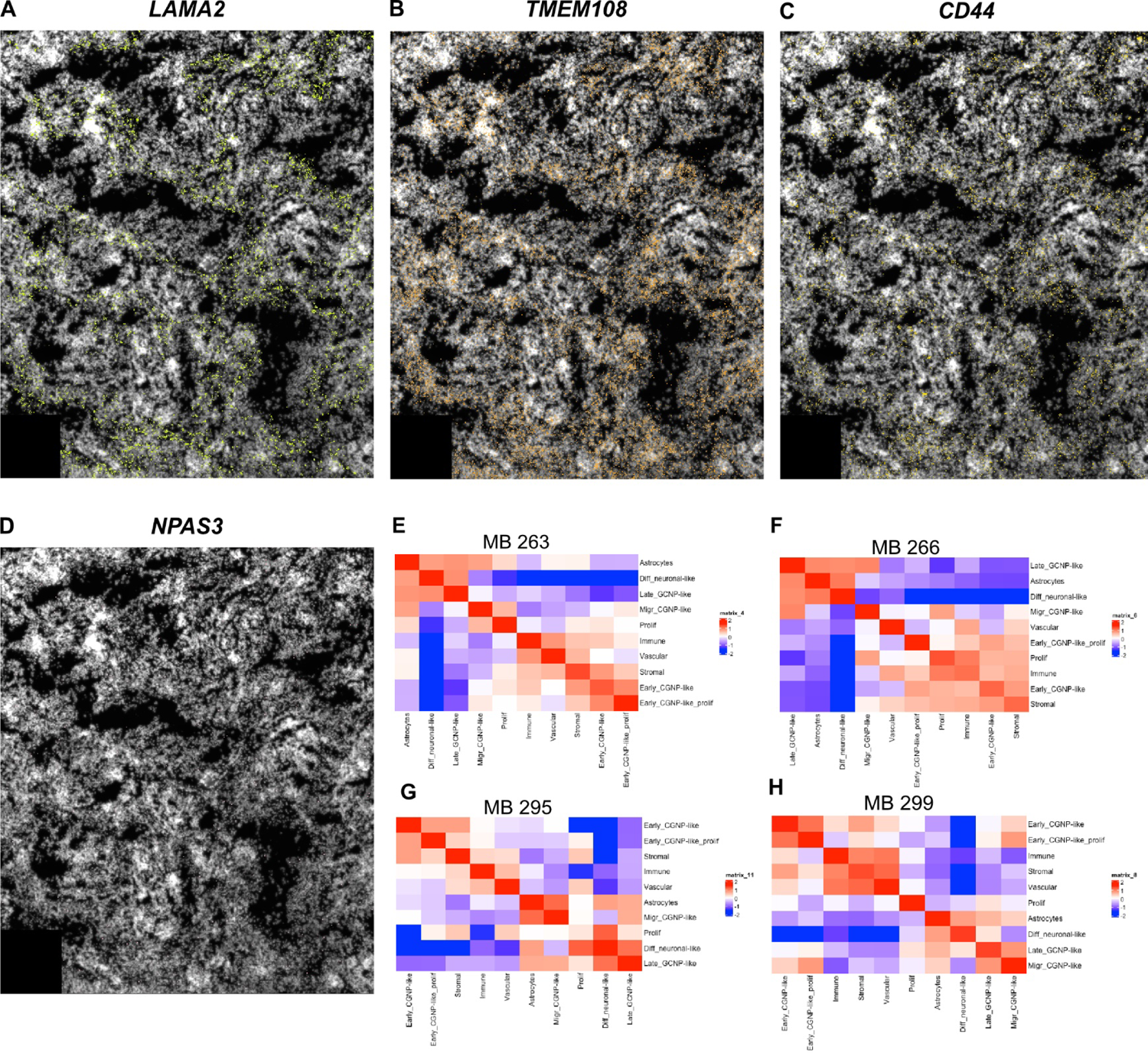
The tumor microenvironment in MBEN differs between its two compartments. Spatial gene expression in MB263 of **A** *LAMA2* (stromal cells, astrocytic cells) **B** *TMEM108* (marker for the internodular compartment), **C** *CD44*, and **D** *NPAS3* (both markers for astrocytic cells) **E - H** Heatmaps summarizing the cell proximity analysis in each case. Cell types which high correlation are more likely to be located next to each other in the tumor microenvironment.

**Suppl. Fig. 12.**
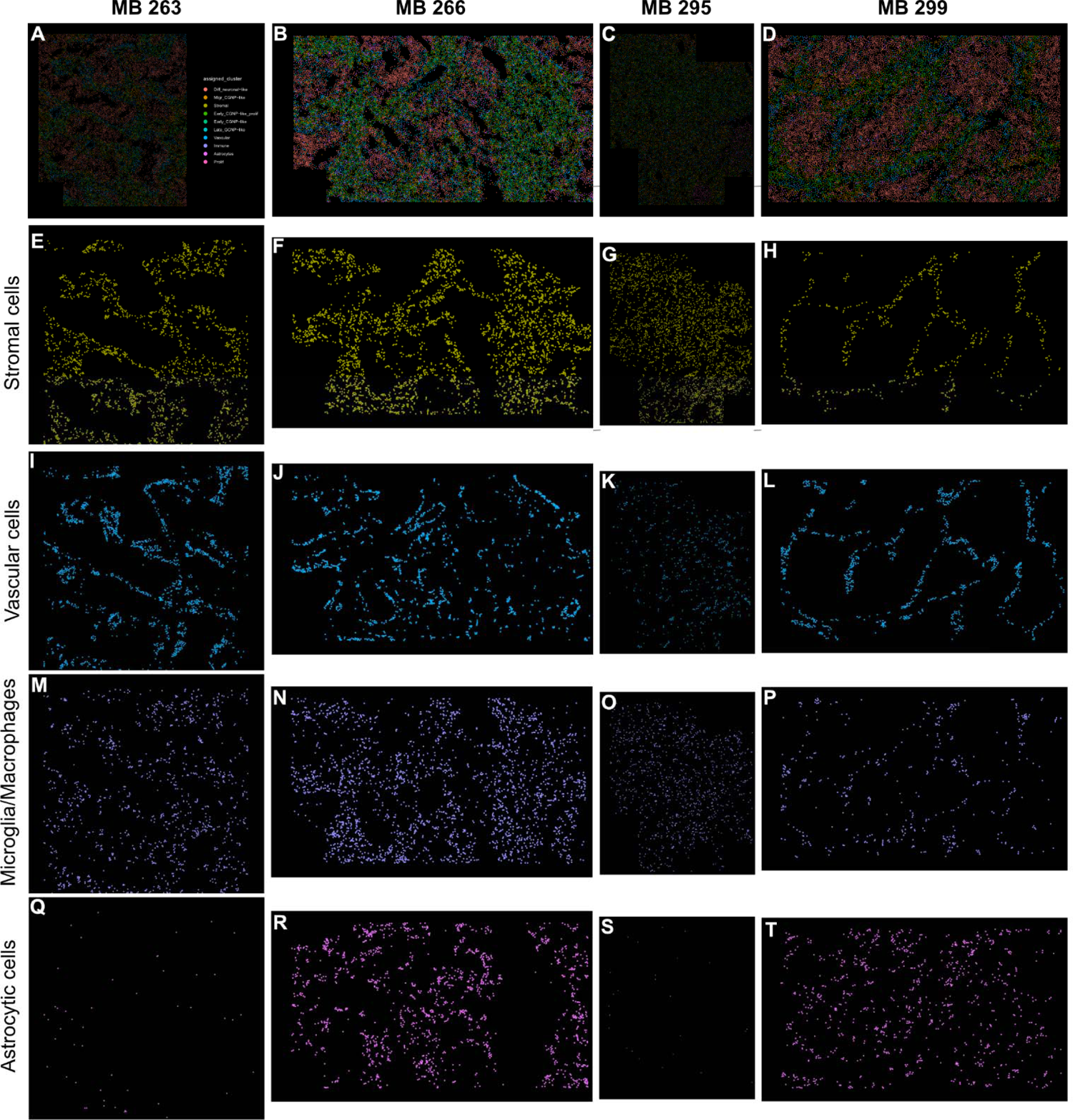
Non-malignant cells in MBEN are enriched in the internodular compartment. **A – D** Scans of four representative MBEN cases with mapping of single cells (legend in A) which recapitulate MBEN histology. **E – H** Mapping of stromal cells. **I – L** Mapping of vascular cells. **M – P** Mapping of microglia. **Q – T** Mapping of non-malignant astrocytes and tumor astrocytes.

